# Wnt-inducible Lrp6-APEX2 Interacting Proteins Identify ESCRT Machinery and Trk-Fused Gene as Components of the Wnt Signaling Pathway

**DOI:** 10.1101/2020.05.03.072579

**Authors:** Gabriele Colozza, Yasaman Jami-Alahmadi, Alyssa Dsouza, Nydia Tejeda-Muñoz, Lauren V. Albrecht, Eric Sosa, James A. Wohlschlegel, Edward M. De Robertis

## Abstract

The canonical Wnt signaling pathway serves as a hub connecting diverse cellular physiological processes, such as β-catenin signaling, differentiation, growth, protein stability, macropinocytosis, and nutrient acquisition in lysosomes. We have proposed that sequestration of β-catenin destruction complex components in multivesicular bodies (MVBs) is required for sustained canonical Wnt signaling. In this study, we investigated the events that follow activation of the canonical Wnt receptor Lrp6 using an APEX2-mediated proximity labeling approach. The Wnt co-receptor Lrp6 was fused to APEX2 and used to biotinylate targets that are recruited near the receptor during Wnt signaling at different time periods. Lrp6 proximity targets were identified by mass spectrometry, and revealed that many components of the ESCRT (Endocytic Sorting Components Required for Transport) machinery interacted with Lrp6 within 5 minutes of Wnt3a treatment. This supports the proposal of a central role of multivesicular endosomes in canonical Wnt signaling. Interestingly, proteomic analyses identified the Trk-fused gene (TFG), previously known to regulate the cell secretory pathway and to be rearranged in thyroid and lung cancers, as being strongly enriched in the proximity of Lrp6. We provide evidence that TFG specifically co-localized with MVBs after Wnt stimulation. TFG depletion with siRNA, or knock-out with CRISPR/Cas9, significantly reduced Wnt/β-catenin signaling in cell culture. *In vivo*, studies in the *Xenopus* system showed that TFG is required for endogenous Wnt-dependent embryonic patterning. The results suggest that the multivesicular endosomal machinery and the novel player TFG have important roles in Wnt signaling.

**Significance:** Wnt/β-catenin signaling is a conserved pathway involved in cell differentiation and in the regulation of many other processes, including cell growth and proliferation, macropinocytosis, and cell metabolism. Endocytosis is required to regulate Wnt signaling, but the precise factors at play are still elusive. Here, we describe a biotin-dependent proximity labeling approach using ascorbate peroxidase-tagged Lrp6, a Wnt co-receptor. Proteomic analysis of biotinylated-enriched targets identified numerous multivesicular endosome proteins that were recruited to the receptor shortly after addition of Wnt protein. Additionally, we identified the protein TFG as one of the strongest interactors with Lrp6. TFG co-localized with Wnt-induced multivesicular endosomes. *Xenopus* embryo assays revealed that TFG is required *in vivo* for canonical Wnt signaling.

## Introduction

The Wnt/β-catenin pathway is an evolutionarily conserved signaling cascade that plays a fundamental role in animal embryonic development, adult stem cell homeostasis, and is dysregulated in a number of human diseases, including cancer (1, 2). In the absence of Wnt stimulation, the scaffold proteins Axin and Adenomatous Polyposis Coli (APC) form a ‘destruction complex’ with Casein Kinase1 (CK1) and Glycogen Synthase Kinase 3β (GSK-3β) which sequentially phosphorylate β-catenin and mark it for ubiquitination and proteasome-mediated degradation (3). Canonical Wnt signal transduction is initiated when a Wnt ligand binds to the seven-pass transmembrane receptor Frizzled (Fzd) and the Low-density lipoprotein receptor-related protein 5/6 (Lrp5/6). The activated co-receptors recruit a number of effectors, including CK1 isoforms (4), the adaptor Dishevelled (Dvl), Axin1 and GSK-3β, to the cytosolic side of the plasma membrane forming a complex known as the signalosome (5). Signaling requires specific protein-protein interactions and the phosphorylation of several components, including the cytoplasmic tail of Lrp6 (6-9). Subsequently, activated Wnt receptor clusters (10) are internalized into multivesicular bodies (MVBs) by ESCRT-driven microautophagy of GSK-3β and other negative Wnt regulators such as Axin, sequestering them away from their cytosolic targets (11, 12). As a result, β-catenin protein is stabilized and translocates into the nucleus, where it binds to T-cell factor/lymphoid enhancer-binding factor (TCF/LEF) DNA-binding proteins to regulate the transcription of Wnt-target genes.

During the 30 years since its discovery (13, 14), the Wnt pathway has been the subject of intense studies, but its molecular mechanisms have not yet been fully elucidated, in particular the role of membrane trafficking in receptor activation (11, 12, 15-17). To dissect the dynamics of Wnt signaling, we decided to use a protein interaction approach to identify additional regulators of the pathway. In recent years, proximity-dependent biotin labeling approaches have emerged as a valuable tool to define functional protein complexes or organelle-specific proteomes (18). Two classes of enzymes can be used in proximity labeling (PL) experiments: the promiscuous *E*. *coli* biotin ligase BirA R118G (BirA*) (19) and its variants TurboID and miniTurboID (20), or the engineered soybean Ascorbate Peroxidase 2 (APEX2) (21). These enzymes attach biotin-derivatives covalently to nearby proteins *in vivo*, while still in their native cellular environment. Biotinylated targets can then be extracted, purified by streptavidin-bead pull-down, and identified by mass spectrometry (MS). The APEX2 peroxidase technology offers several advantages: faster kinetics, robust biotinylation activity, and the possibility to activate the labeling at specific moments by adding H_2_O_2_ to living cells (21, 22). APEX2 requires the use of biotin-phenol (also known as biotin-tyramide), which in combination with H_2_O_2_ is converted to a highly reactive, short-lived (<1 msec) radical with a labeling radius of 20 nm. The high spatial and temporal specificity provided by APEX2 has allowed the study of the proteome of organelles or subcellular compartments difficult to isolate, such as the mitochondrial intermembrane space and endoplasmic reticulum (ER) membrane (23), and to study G-protein-coupled receptor (GPCR) signaling in great detail (24, 25).

Here, we describe the results obtained with a Human Embryonic Kidney 293T (HEK293T) cell line stably expressing a chimeric Lrp6-APEX2 receptor. Our proteomic data indicate that in presence of Wnt3a, the Lrp6-interactome was enriched in components of the ESCRT machinery that forms MVBs, providing independent support to our previous findings that endocytosis is at the core of Wnt pathway intracellular regulation (11, 15, 16). We also discovered that Tropomyosin-receptor kinase fused gene (Trk-fused gene, TFG), a protein involved in several pathologies including cancer (26), was a highly enriched target of Wnt-activated Lrp6-APEX2 and localized to endosomal vesicles. Cell culture knock-out assays and *in vivo* experiments using *Xenopus laevis* embryos revealed an important role for TFG in Wnt/β-catenin signaling and antero-posterior (AP) patterning.

## Results

### Lrp6-APEX2 Receptor Fusion is Active in Wnt Signaling

To study the protein complexes that are recruited in proximity of the receptor complex during Wnt signaling, we appended APEX2 (21) to the cytoplasmic tail of the Lrp6 receptor (Fig. 1 *A* and *B*). The chimeric receptor maintained functionality, since mRNA overexpression could rescue the ventralized phenotype caused by maternal depletion of *Xenopus* Lrp6 (27) in frog embryos (Fig. 1 *C*-*E*), and induced axis duplication in a ligand dependent manner when overexpressed in a ventral blastomere (*SI Appendix*, Fig. S1 *A*-*G*), as is the case with the wild type receptor (6). Importantly, Lrp6-APEX2 also restored the response to Wnt ligands by Lrp5/6 double knock-out HEK293 cells (28), corroborating its functionality (*SI Appendix*, Fig. S1*H*). Next, we established a HEK293T stable clonal line expressing the Lrp6-APEX2 construct, which included the GFP-tagged puromycin selectable marker eGFP-PAC after an internal ribosome entry site (IRES) (Fig. 1*A*). Lrp6-APEX2 was robustly expressed (*SI Appendix*, Fig. S1*I*), and was properly trafficked to the plasma membrane (Fig. 1 *F*-*H*). Treatment with Wnt3a conditioned medium translocated Flag-tagged Lrp6-APEX2 into cytoplasmic vesicles within 30 minutes (compare Fig. 1 *I* to *J*).

**Fig. 1.**
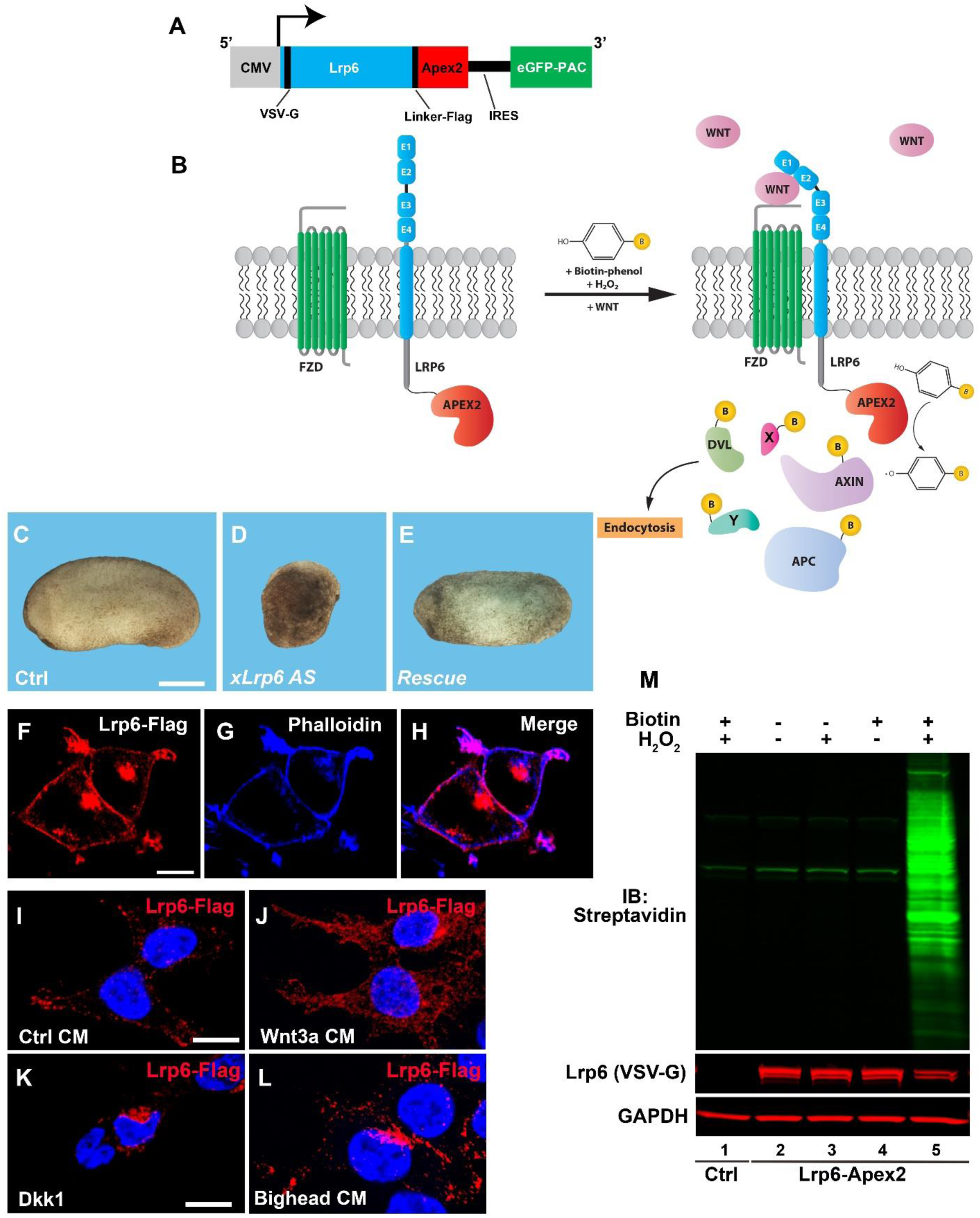
The Lrp6-APEX2 fusion protein receptor retains signal transducing activity, proper subcellular localization, and biotinylates cellular proteins. (*A*) Schematic diagram of the Lrp6-APEX2 construct used in this study to establish stable cell line. The human Lrp6 contains an N-terminal VSV-G and a linker Flag-tag followed by the APEX2 enzyme. The construct also contains an internal ribosomal entry site (IRES) followed by a GFP-tagged Puromycin N-acetyltransferase (PAC) used for selection of permanently transfected cell clones. (*B*) Diagram depicting the biotinylation of Lrp6-APEX2 proximal targets. During the Wnt signal, in the presence of Biotin-phenol (BP) and H_2_O_2_, proteins recruited to the receptor are biotinylated by the APEX2 peroxidase, including known Wnt targets (Dvl, Axin, APC, etc.) and novel targets (X, Y). (*C-E*) control embryos from oocyte-host transfer experiments show normal development (100%, n=24), as compared to oocytes depleted with a phosphorothioate (*) antisense DNA oligo (27) targeting *Xenopus* Lrp6 (T*C*G*AGGCTGATCCAG*C*T*C), which develop as ventralized embryos (75%, n=22). Co-injection of 300 pg of Lrp6-APEX2 mRNA into oocytes completely rescued axis formation (60%, n=17), indicating that the fusion protein is fully active; additional experiments supporting this conclusion are shown in *SI Appendix*, Fig. S2. Scale bar represents 500 µm. (*F-H*) Lrp6-APEX2 is trafficked to the plasma membrane in stably transfected HEK293T cells. Lrp6 was detected through its Flag-tag and Phalloidin, which stains cortical F-Actin, confirmed cell surface localization of Lrp6. Note that a lower concentration of Triton X-100 detergent was used (0.05%), to improve membrane staining (see materials and methods). Scale bar represents 10 µm. (*I-L*) Lrp6-APEX2 changes subcellular localization by Wnt, Dkk1 and Bighead treatments. When treated with control conditioned medium (CM), Lrp6 is located at the plasma membrane and a small number of intracellular vesicles. Treatment with Wnt3a CM for 30 minutes increases the number of intracellular vesicles containing Lrp6, indicating endocytosis of the receptor. Treatment with Dkk1 protein (200 ng/ml) or Bighead CM induced relocation of Lrp6 to the juxtanuclear bay area where lysosomes are located. Scale bars represent 10 µm. (M) western blot stained with Streptavidin-IRDye 800 (in green) showing controls indicating that Lrp6-APEX2 biotinylates proximal proteins only in the presence of both BP and H_2_O_2_ (compare lane 5 with 2-4). Negative control cells not expressing the APEX2 peroxidase had no biotinylation even in presence of BP/H_2_O_2_ (lane 1). Note the presence of three Streptavidin-positive bands in all lanes, which correspond to endogenous biotinylated carboxylases. The VSV-G immunoblot confirmed that Lrp6 was expressed in these cells, while Gapdh served as a loading control.

Wnt activation is known to induce endocytosis of the receptor complex (11, 12). We confirmed that Lrp6 endocytosis was observed in SW480, a colon cancer cell line in which the absence of APC triggered, and APC reconstitution inhibited, Wnt signaling and Lrp6 endocytosis (*SI Appendix*, Movie S1); this is in agreement with recent results showing that APC is a major regulator of endocytosis (16, 29). As expected for regulators of Lrp6 endocytosis, the Wnt antagonists Dkk1 (30) or Bighead (31) induced endocytosis of Lrp6-APEX2 into large vesicle clusters in the cytoplasm next to the nuclear bay region, where the lysosomal system is located (Fig. 1 *K* and *L*). Next, we assessed the activity of the APEX2 enzyme. The western blot in Fig. 1*M* includes several control treatments showing that APEX2-mediated biotinylation only occurred in the presence of both biotin-phenol and H_2_O_2_. This activity could also be detected by Streptavidin immunofluorescence (*SI Appendix*, Fig. S1 *J*-*O*). Altogether, our data show that the chimeric receptor Lrp6-APEX2 can replace the wild type receptor and can be used for biotin-labeling studies.

Compared to the parental HEK293T cell line, Lrp6-APEX2 HEK293T cells showed elevated levels of phosphorylated Dvl2 (pDvl2) (*SI Appendix*, Fig. S2*A*); however, this Dvl2 phosphorylation was blocked upon preincubation with the Porcupine inhibitor IWP-2 (32), indicating a higher sensitivity to an unknown endogenous Wnt present in HEK293T cells (*SI Appendix* Fig. S2*B*). In the presence of IWP-2, high levels of phospho-Dvl2, active (*i*. *e*., non-phosphorylated) β-catenin and phospho-Lrp6 (pLrp6) were only observed when cells were exposed to Wnt3a conditioned medium and, interestingly, this response was increased by R-Spondin 1 and 2 (*SI Appendix* Fig. S2*C*). Thus, our Lrp6-APEX2 cells are entirely dependent on exogenous Wnt extracellular ligands, providing a new and sensitive system to explore the interactions of the Lrp6 receptor during Wnt signaling.

### Wnt Induces Lrp6-APEX2 Biotinylation of Components of the ESCRT Machinery

Lrp6-APEX2 HEK293T cells (preincubated with Porcupine inhibitor to eliminate endogenous Wnts) were treated for 30 minutes with biotin-phenol in T225 flasks (Fig. 2*A*). Next, duplicate samples were treated with either control or Wnt3a conditioned medium containing biotin-phenol (fortified by the addition of 50 ng/ml R-Spo1 and 2 which were found to strongly increase Wnt3a responsiveness; *SI Appendix*, Fig. S2*D*). Two time points were analyzed, 5 and 20 min. The labeling reaction was triggered by a 1-minute incubation with H_2_O_2_, followed by cell lysis and pull-down of biotinylated proteins (Fig. 2*A*). Each treatment included biological duplicates. Biotinylation and β-catenin stabilization were confirmed by western blot in each sample (*SI Appendix*, Fig. S2*E*). Affinity-purified samples were then processed for tandem mass spectrometry (MS/MS) of tryptic peptides (33, 34) as described in Materials and Methods. All original datasets generated in this study are publicly available at the MassIVE public repository resource (accession number MSV000084335).

**Fig. 2.**
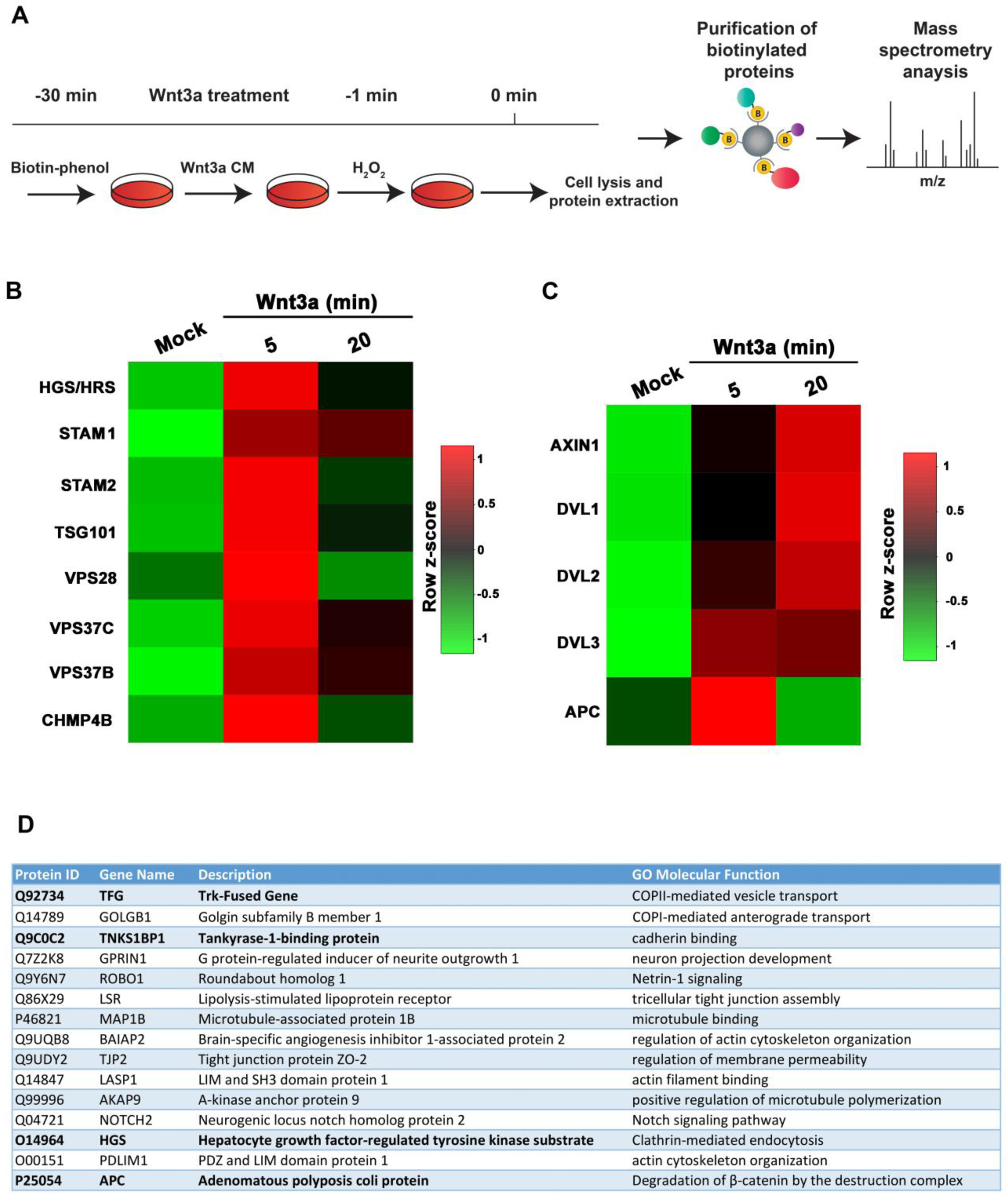
Analysis of Lrp6-APEX2 proximal targets reveal rapid Wnt3a-induced interaction between Lrp6 and ESCRT proteins. (*A*) Schematic of the biotinylation proteomic experiments reported in this study. (*B*) Heatmap of Lrp6-APEX2 proximity labeled ESCRT proteins using normalized intensities at 5 and 20 minute time points, showing the horizontal row z-scores of proteins over time. Mock indicates cells treated with control conditioned medium from cells not expressing Wnt3a. Row z-scores were calculated from average intensities of proteins. Average intensity values were obtained from biological duplicates. Highly biotinylated proteins are shown in red and lower ones in green. Note the enrichment in ESCRT proteins particularly after 5 minutes of Wnt3a treatment. (*C*) Heatmap of Lrp6-APEX2 proximity labeled known canonical Wnt signaling pathway target proteins using normalized intensities at 5- and 20-min time points, showing the horizontal row z-scores of proteins over time. Average intensity values were obtained from biological duplicates. Highly biotinylated proteins are shown in red and lower ones in green. Note the interaction between Lrp6-APEX2 and Dvl 1-3 and Axin proteins after 20 minutes of Wnt3a treatment while APC peaks at 5 min; these interactions support the specificity of our Wnt3a signaling experiments. (*D*) The top 15 biotinylated targets (ranked according to their spectral counts) from Dataset S1 Tab 1. Note the presence of the ESCRT-0 protein Hrs/Hgs, the Wnt inhibitor APC and Actin remodeling proteins such as BAIAP2, LASP1 and PDLIM1. Unexpectedly, the Trk-fused gene (TFG) was the most enriched protein in this list.

Principal Component Analysis (PCA) showed that each replicate clustered according to the type of treatment, validating the reproducibility of the assay (*SI Appendix*, Fig. S2*F*). Analysis of the peptide data generated a list of over 4,000 putative biotinylated proteins (all of which are listed in Tab 1 of Dataset S1). Of these, over 2,000 could be accurately quantified using MS1-based label free quantitation (Dataset S1, Tab 2), as explained in Materials and Methods. Of those 2,000-plus proteins, 217 and 237 proteins were significantly enriched (p-Value <0.05) in the 20 and 5 minute Wnt3a samples, respectively, compared to no biotin controls (Dataset S1, Tab 3). Amongst the most highly enriched proteins was the Trk-Fused Gene (TFG). The full list of identified proteins (including their spectral counts) that interact with Lrp6-APEX2 after 5 or 20 min of Wnt3a treatment is shown in Tab 1 of Dataset S1 and represents a rich resource to search for downstream interactions of the Lrp6 receptor during Wnt signaling.

Gene Ontology analyses using the Panther database revealed that the top hit induced by Wnt3a conditioned medium was the ESCRT machinery required for MVB formation. Heatmaps derived from the normalized list of proteins showed a clear involvement of the ESCRT machinery, with a marked enrichment of Hrs, Stam1/2, Chmp4, Vps28, Vps35b, Vps35c and Tsg101 among others, which was strongest after 5 minutes of Wnt3a treatment (Fig. 2*B*). Confirming that the Lrp6-APEX2 interactions were Wnt-induced, known targets of the canonical Wnt pathway, such as APC, Dvl 1-3 and Axin and were biotinylated upon Wnt3a treatment, although in this case the interactions at 20 min were maximal, except for APC which was maximal at 5 min (Fig. 2*C*). APC was located in positon 15 and Dvl2 in position 38, the complete list is available in Dataset S1, Tab 1. Proteins involved in actin cytoskeleton remodeling were also prominent among Wnt-inducible interactions. Since canonical Wnt signaling triggers macropinocytosis through the actin contractile machinery, this might be indicative of a role of Lrp6 in plasma membrane macropinocytosis (16, 37). Fig. 2*D* shows a partial list including the 15-most differentially enriched biotinylated proteins upon a 20 min Wnt3a treatment, which included TFG, Tankyrase-1 binding protein, Hrs/Hgs and APC.

We also carried out proteomic analysis of a single sample treated for 1 hour with Wnt3a conditioned medium (CM) or CM from control cells not expressing Wnt3a, designated Mock CM (Dataset S1, Tab 4). This longer time point (which did not include the use of R-spondin 1 and 2) revealed that the top three Lrp6-APEX2 interactors were α-Spectrin, β-Spectrin and their ancestor α-Actinin (from which spectrins evolved) (36). These proteins form a meshwork at the plasma membrane (together with protein band 4.1) that serves to link actin filaments to the plasma membrane (36). The continuous involvement of the actin cytoskeletal machinery is consistent with a sustained role for nutrient acquisition via macropinocytosis in canonical Wnt signaling (16, 37). We also performed MS analysis on whole protein extracts derived from the 1 hour Wnt3a treatment, before any streptavidin-based enrichment, confirming the stabilization of β-catenin by Wnt (Dataset S1, Tab 5).

We next analyzed more deeply this global proteomic data using the Metascape online software. Wnt3a-induced biotinylated proteins were ranked for enrichment over Mock control CM samples. The lists of the top 150 proteins were analyzed for Gene Ontology groups at 5 and 20 minutes and 1 hour, as shown in *SI Appendix*, Fig. S3*A-C*. After 5 min endocytosis was the top hit, followed by small GTPase-mediated signal transduction. Interestingly, Kras was notable among Wnt-induced Lrp6 interacting proteins, both after 5 and 20 min. This is important because Kras is a major regulator of many metabolic processes, including actin-driven macropinocytosis. In the 20 minute sample, the top GO score was signaling by Wnt, as might have been expected. After 1 hour of Wnt, actin cytoskeleton organization and membrane trafficking were pre-eminent.

In summary, our searchable list of Lrp6-APEX2 biotinylated targets (Dataset S1, Tab 1 and Tab 4) offers the scientific community a valuable roadmap for the discovery of novel proteins involved in Wnt signaling and endocytosis. Importantly, it independently confirms our earlier proposals (11) that the ESCRT machinery and endocytosis (15, 16) participate in Wnt signaling.

### TFG Localizes to Endocytic Vesicles and Is Required for Wnt Signaling

Unexpectedly, one of the most highly enriched proteins upon 20 minutes of Wnt3a treatment was TFG (Fig. 2*D*), a protein containing multiple domains (*SI Appendix*, Fig. S4*A*). N-terminal in-frame fusions of TFG to other genes are found in thyroid and lung cancers (26, 38, 39). Notably, immunostaining of untransfected HeLa cells revealed endogenous TFG puncta that co-localized with Hrs (Fig. 3 *A*-*C*), an ESCRT-0 protein that is required for Wnt signaling (11). Hrs puncta, as well as their co-localization with TFG, were increased by Wnt3a after only 5 min of treatment (Fig. 3*D-F*, quantified in Fig. 3*G* and *H*). Co-localization of TFG and Hrs was confirmed by co-transfection of TFG-flag and Hrs-GFP constructs, with both proteins enriched in the periphery of endocytic vesicles (*SI Appendix*, Fig. S4 *B*-*D*). Co-immunostaining revealed close association between Lrp6-APEX2 and TFG even in the absence of Wnt (*SI Appendix*, Fig. S4 *E*-*G*). To investigate the function of TFG during Wnt signaling, we performed knock-down experiments in cultured cells. As shown in Fig. 3*I*, transfection of siRNA against TFG for two days decreased signaling by the Wnt/β-catenin-driven BAR luciferase reporter by over 50%, indicating that TFG is required for canonical Wnt signaling in cultured cells.

**Fig. 3.**
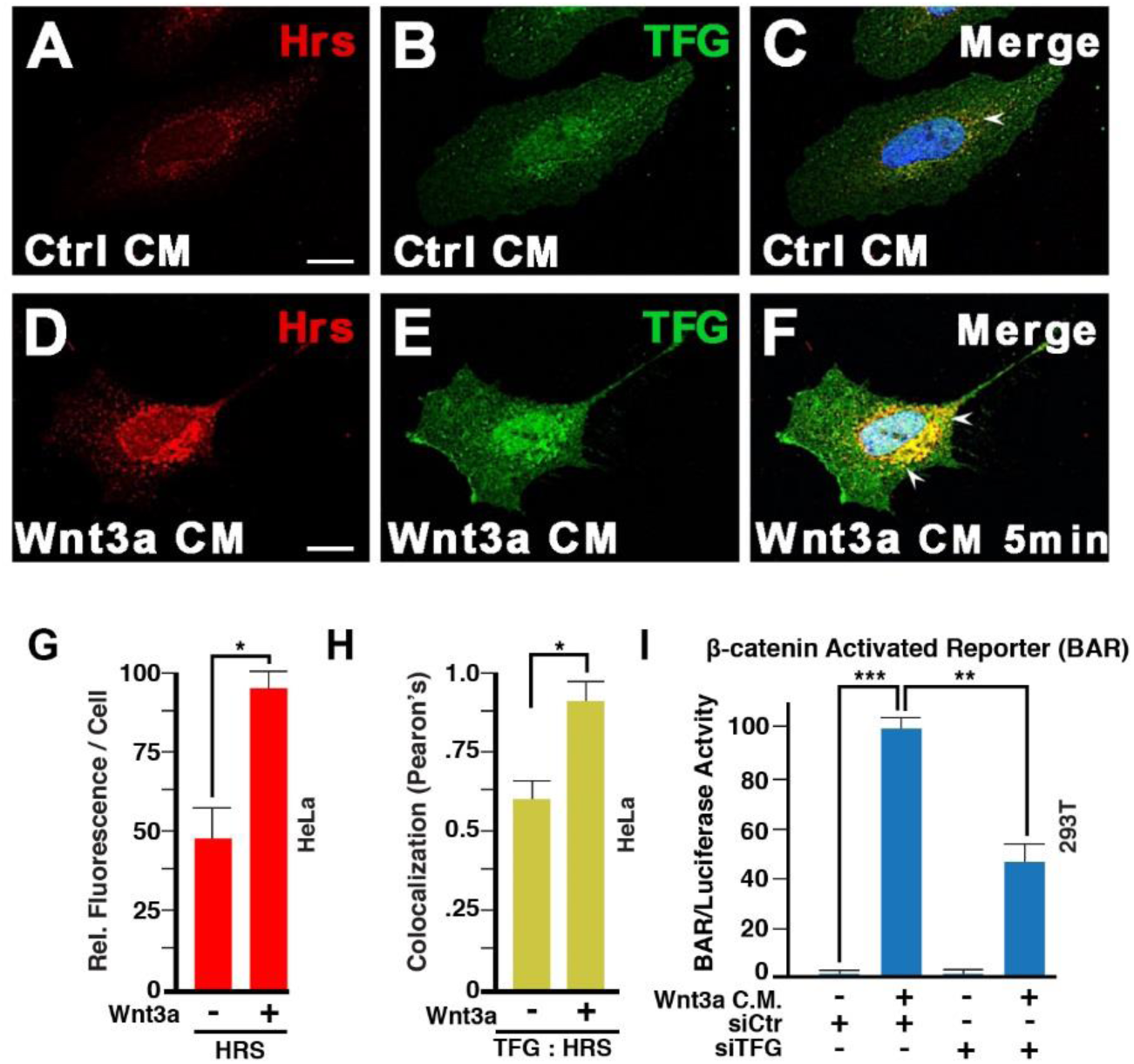
TFG co-localizes with the ESCRT-0 protein Hrs/Hgs and is required for Wnt signaling. (*A-F*) Immunostaining on HeLa cells for endogenous Hrs and TFG. Weak co-localization was observed in absence of Wnt signaling (arrow in panel C). However, 5 minutes of Wnt3a treatment was sufficient to strikingly increase the number of endocytic vesicles containing both Hrs and TFG (arrows). Scale bar=10 µm. (*G*) Wnt signaling (5 min) increased the number of Hrs positive vesicles quantified as fluorescence per cell; this supports the rapid formation of MVBs in Wnt signaling. (H) Wnt3a increased co-localization of TFG puncta with the Hrs MVB marker by Pearson’s correlation coefficient. (I) TFG knock-down by siRNA reduced Wnt signaling, as assessed by β-catenin Activated Reporter (BAR/Renilla) Luciferase assay in HEK293T cells.

To further demonstrate the role of TFG in regulating Wnt signaling, we used the CRISPR/Cas9 (Clustered regularly interspaced short palindromic repeats/CRISPR-associated 9) gene editing technology (40-42) to achieve full TFG knock-out. A guide RNA (sgRNA) targeting a sequence proximal to TFG start codon (Fig. 4*A*) was subcloned into a Cas9 expression vector and transfected into HEK293T cells harboring BAR Luciferase/Renilla reporters (15). Western blot analyses confirmed complete depletion of TFG protein in a knock-out clonal cell line derived by limiting dilution (Fig. 4*B*). This knock-out line had two independent frameshift mutations caused by insertion/deletion (indel). Both mutations introduced an early termination codon, resulting in truncated TFG proteins of approximately 90 amino acids (*SI Appendix*, Fig. S5). Next, we studied Wnt signaling in TFG knock-out cells. In wild-type (WT) cells, treatment with Wnt3a CM for 3 hours induced a substantial increase in β-catenin immunostaining in WT cells (Fig. 4*C, E*), whereas TFG KO cells showed some increase in β-catenin upon Wnt3a treatment but at much reduced extent when compared to WT cells (Fig. 4, compare panels *E* and *F*). This suggested that TFG was required for maximal Wnt-induced β-catenin stabilization. Moreover, deletion of TFG reduced Wnt-driven BAR-luciferase reporter activity by over 50% compared to WT cells (Fig. 4*G*), in agreement with our previous observations with TFG siRNA. An even stronger reduction in β-catenin signaling (80%) was observed when we used CHIR99021, a specific GSK3 inhibitor, instead of Wnt3a in TFG mutant cells (Fig. 4*H*). While a requirement of TFG for GSK3 inhibition signaling was unexpected, it might be explained by our recent findings that CHIR99021 triggers massive macropinocytosis (Albrecht et al, submitted); perhaps membrane trafficking, in which TFG is involved, is required for β-catenin stabilization. Taken together, the results suggest that the Lrp6 interacting protein TFG is indeed a novel component of the canonical Wnt signaling pathway.

**Fig. 4.**
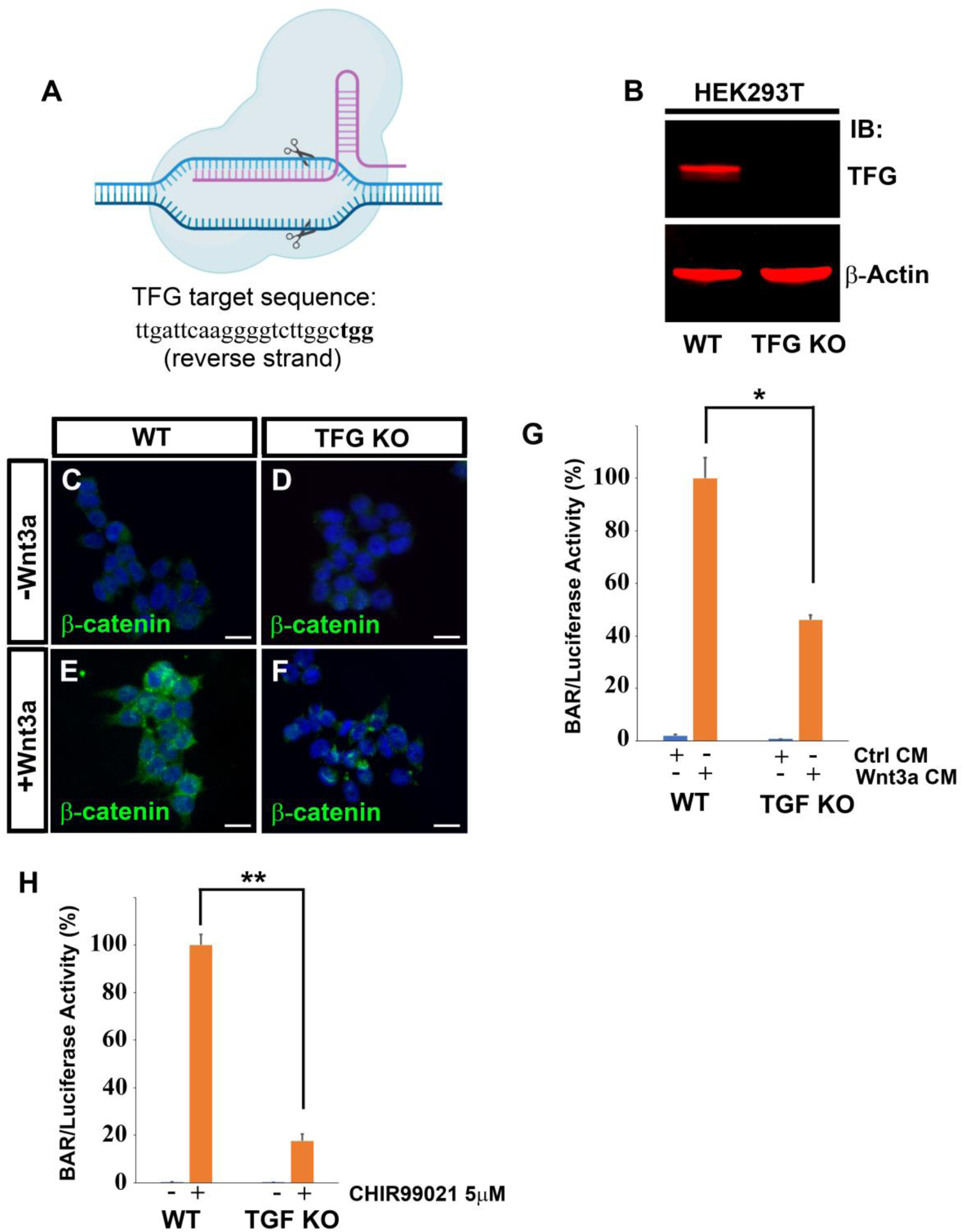
CRISPR-Cas9 mediated TFG knock-out inhibits Wnt-dependent β-catenin stabilization and Wnt-induced reporter activity. (*A*) Schematic diagram of Cas9 cleaving the TFG genomic DNA target sequence. The target sequence (which is on the reverse strand) is shown in 5’ to 3’ orientation (left to right); the PAM (protospacer adjacent motif) sequence is in bold. (*B*) Western blot of HEK293T cells confirming complete elimination of TFG protein in Cas9 knock-out cells. β-actin was used as loading control. (*C-F*) Immunostaining for β-catenin on WT or TFG KO cells. Cells were treated with control conditioned medium or with Wnt3a conditioned medium for 3 hours before immunostaining. Note that TFG knock-out decreases Wnt3a-induced β-catenin accumulation; DAPI was used for nuclear counter-staining. Scale bars represent 20 µm. (*G*) Cas9-mediated TFG knock-out reduces response to Wnt3a by 50% in HEK293T BAR/Renilla β-catenin reporter cells. Cells were treated with Wnt3a or control medium for 16 hours before luciferase analysis. (*H*) Luciferase assay of HEK293T BAR reporter cells treated with the GSK3 inhibitor CHIR99021 at 5 µM concentration for 16 hours before being processed for luciferase assay. Note that TFG KO causes an 80% reduction of β-catenin luciferase activity in response to the GSK3 inhibition. Error bars represent standard deviation from triplicate experiments. Statistical significance was calculated with a paired 2-tailed t-Student test. * = P < 0.05; ** = P < 0.01.

### TGF is Required for Wnt Signaling in *Xenopus* Embryos

To address the developmental role of TFG *in vivo*, we turned to *Xenopus* embryos as a model of choice. RT-PCR data showed that TFG was a maternal gene and its expression remained uniform until late stages of development (*SI Appendix*, Fig. S4*H*). Unlike Chordin, a prototypical Spemann organizer gene (43), TFG did not show dorso-ventral polarity (*SI Appendix*, Fig. S4*I*). *In situ* hybridization showed that TFG was expressed in the animal pole of early cleavage embryos, and was later strongly expressed in the cement gland, as reported by others (*SI Appendix*, Fig. S4 *J*-*N*) (44), in addition to staining in the notochord as shown here (*SI Appendix*, Fig. S4*O*).

To perform loss of function studies *in vivo*, we designed a morpholino antisense oligo (45) which targeted both *Xenopus laevis* TFG homeologs (Fig. 5*A*) and efficiently blocked the translation of microinjected *Xenopus* flag-tagged *TFG* mRNA but not human *TFG* mRNA differing in the target sequence (*SI Appendix*, Fig. S6*A*). Embryos microinjected with TFG morpholino displayed a strong anteriorization, with an enlargement of the cement gland and head (Fig. 5 *B* and *C*, and *SI Appendix*, Fig. S6*B*). Head enlargement was confirmed by *in situ* hybridization using the anterior marker *Otx2* (*SI Appendix*, Fig. S6 *C* and *D*). Co-injection of human Flag-tagged *TFG* mRNA completely rescued the anteriorizing effect, demonstrating that the phenotype of the TFG morpholino oligo was specific (Fig. 5*D*). Overexpression of human or *Xenopus* TFG mRNAs were without phenotypic effects on their own. However, human TGF lacking the MO target sequences was able to rescue the large head structures caused by TFG knockdown (Fig. 5*D*). The phenotype of TFG knock-down was very similar to that caused by the overexpression of known Wnt antagonists, such as Dkk1 (Fig. 5*E*), indicating that TFG is required for Wnt signaling *in vivo*. In line with this, TFG loss of function in one half of the embryo (marked by co-injection of nuclear *LacZ* mRNA) caused a marked decrease in the expression of the Wnt target gene *Engrailed-2* (Fig. 5 *F* and *G*), while expression of the hindbrain marker *Krox20* was still present although shifted posteriorly. The pan-neural marker *Sox2* was expanded by TFG morpholino, an effect also observed upon Wnt inhibition (46), and was rescued by co-injection with human *TFG* mRNA (Fig. 5 *H*-*J*).

**Fig. 5.**
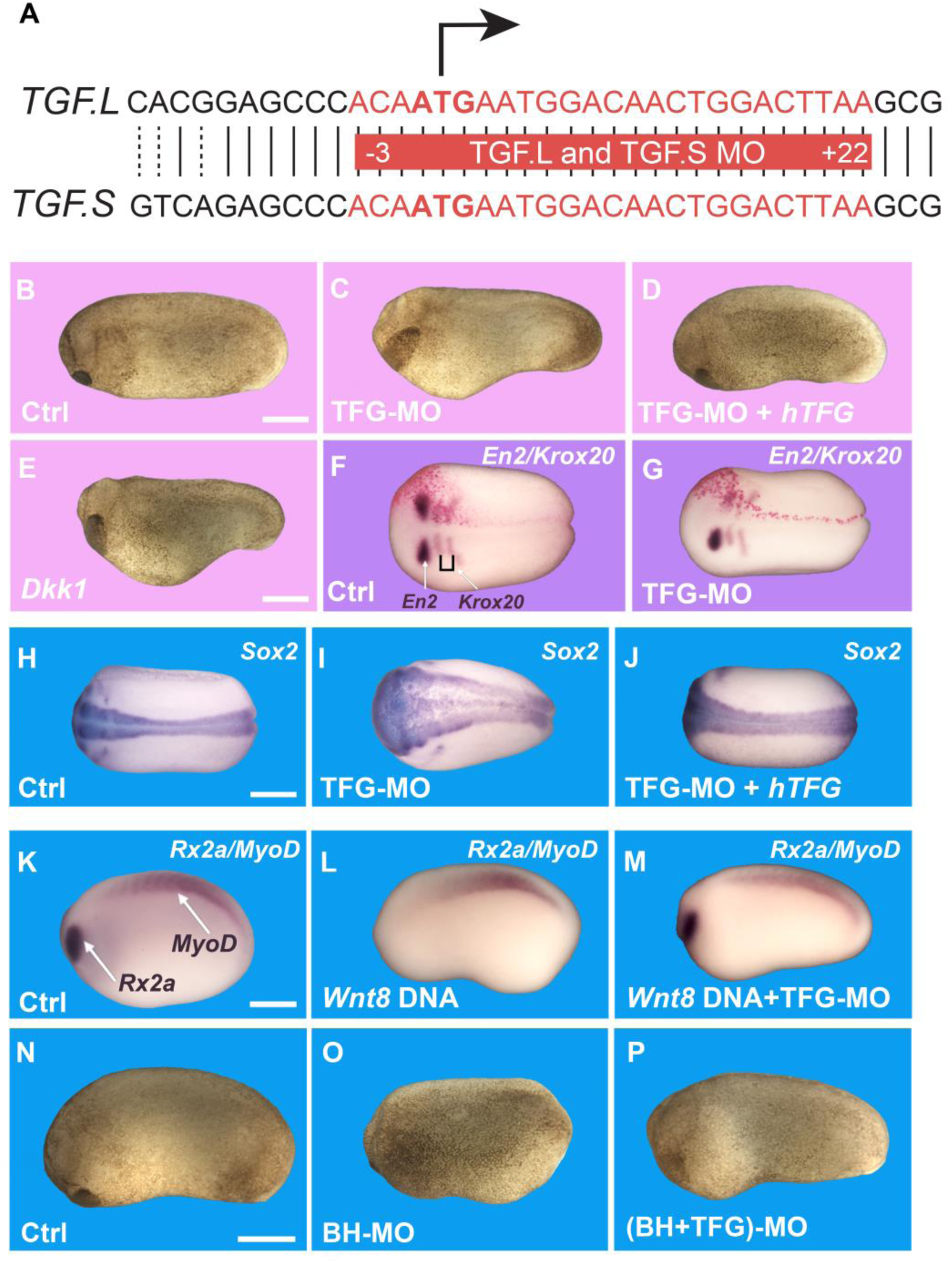
TFG regulates antero-posterior (A-P) patterning in the *Xenopus* embryo and is required for Wnt activity. (*A*) Diagram showing the nucleotide sequences recognized by the translation-blocking TFG morpholino (MO). The target sequence is the same for both *Xenopus* TFG.L and TFG.S homeologs present in the sub-tetraploid *Xenopus laevis* genome, and is shown in red, spanning the ATG initiation codon. (*B*) Embryos injected with control morpholino (Co-MO) were fixed at stage 24, showing normal development (92%, n=127). (*C*) Embryos injected with 36 ng of TFG-MO into the two dorso-animal blastomeres at the 8-cell stage showed severe enlargement of the head region (65%, n=196). *(D)* Normal antero-posterior development was rescued when 400 pg/embryo of human Flag-tagged *TFG* mRNA was co-injected with the morpholino (77%, n=52). (*E*) Embryos injected 4 times at the 4-cell stage with 50 pg of mRNA encoding the Wnt antagonist Dkk1 showed a phenotype similar to TFG loss of function (100%, n=35). (*F, G*) Embryos injected unilaterally at the 2-cell stage with 18 ng of Co-MO or TFG-MO, together with 200 pg of nuclear *LacZ* mRNA, fixed at stage 15, and processed for *in situ* hybridization for *Engrailed-2* (*En2*, a midbrain-hindbrain marker and a known canonical Wnt target) and *Krox20*, a hindbrain marker. LacZ lineage tracing (in red) shows the injected side. Note that TFG-MO reduces *En2* expression and shifts *Krox20* posteriorly in 65% of the embryos (n=20), an effect not observed in embryos injected with scrambled control MO (n=33). (*H* and *I*) *In situ* hybridization for the pan-neural marker *Sox2* at stage 20 showing enlargement of the neural plate following injection 4 times at the 2-cell stage with a total of 72 ng of TFG-MO per embryo (70%, n=65), compared to the Ctrl-MO embryos (n=35). (*J*) Co-injection with 800 pg of *hTFG* mRNA rescued the neural plate expansion caused by TFG-MO (78%, n=23). (*K*-*M*) Experiment showing that the posteriorizing effect of pCSKA-Wnt8 DNA requires TFG activity. Control embryos at stage 22 showed normal eye (*Rx2a*) and somite (*MyoD*) development (n=18), while embryos injected 4 times at the 4 cell-stage with 32 pg of *xWnt8* DNA together with 72 ng of scrambled Ctrl-MO showed reduced or no *Rx2a* expression, but maintained *MyoD* expression (66%, n=18). However, embryos injected with the same amount of *Wnt8* DNA together with 72 ng of TFG-MO showed rescue of *Rx2a* expression (84%, n=32), indicating that TGF is required for xWnt8 signaling. (*N*-*P*) embryos injected with Bighead morpholino (BH-MO) showed strong reduction of head and cement gland development (71%, n=21) when compared to control embryos (n=37). This is explained by an increase in plasma membrane Lrp6 and hence increased endogenous Wnt signaling (31). Note that co-injection of TFG-MO rescued normal head development in Bighead MO embryos (90%). Scale bars in *B*-*P* represent 500 µm.

Next, we injected xWnt8 DNA into *Xenopus* embryos that, as expected (47), caused the inhibition of anterior structures such as eyes (Fig. 5 *K* and *L*) and cement gland (*SI Appendix*, Fig. S6 *E* and *F*). However, injection of TFG-MO together with Wnt8 DNA rescued eye and cement gland development (Fig. 5 *M* and *SI Appendix*, Fig. S6*E*-*G*), indicating that TFG is required for Wnt activity *in vivo*. To show that TGF-MO is required for endogenous late Wnt signaling we used knock-down of the Wnt antagonist Bighead, which is known to result in an increase of Wnt activity (31). We found that while Bighead-MO caused loss of anterior structures in *Xenopus* embryos, this effect could be rescued by co-injection of TFG-MO (Fig. 5*N*-*P*), indicating that TFG functions downstream of Lrp6. Taken together, the results indicate that the Lrp6-APEX2 interacting protein TFG is required for Wnt/β-catenin signaling in *Xenopus* development.

## Discussion

### The ESCRT Machinery and Wnt Signaling

In the present study, we used biotin-dependent proximity labeling to study the early molecular interactions following Wnt/β-catenin activation. Proteomic analyses generated a searchable list of Lrp6 interactors that is provided in Dataset S1 Tab 1, and can be mined by the research community to identify additional Wnt regulators. Importantly, our data showed that ESCRT proteins such as Hrs/Hgs, Stam1/2, CHMP4, Vps28, Vps37 and Tsg101 were recruited close to the Lrp6 Wnt receptor soon after Wnt stimulation (5 min) and were still significantly enriched 20 min later. In agreement with this, our immunostaining on cultured cells showed that 5 min of Wnt3a treatment was sufficient to significantly increase the number of Hrs-containing endocytic vesicles. Thus, the results presented here strongly support the view that multivesicular endosome formation is a key downstream step in Wnt signaling (11, 12), as indicated in the model shown in Fig. 6.

**Fig. 6.**
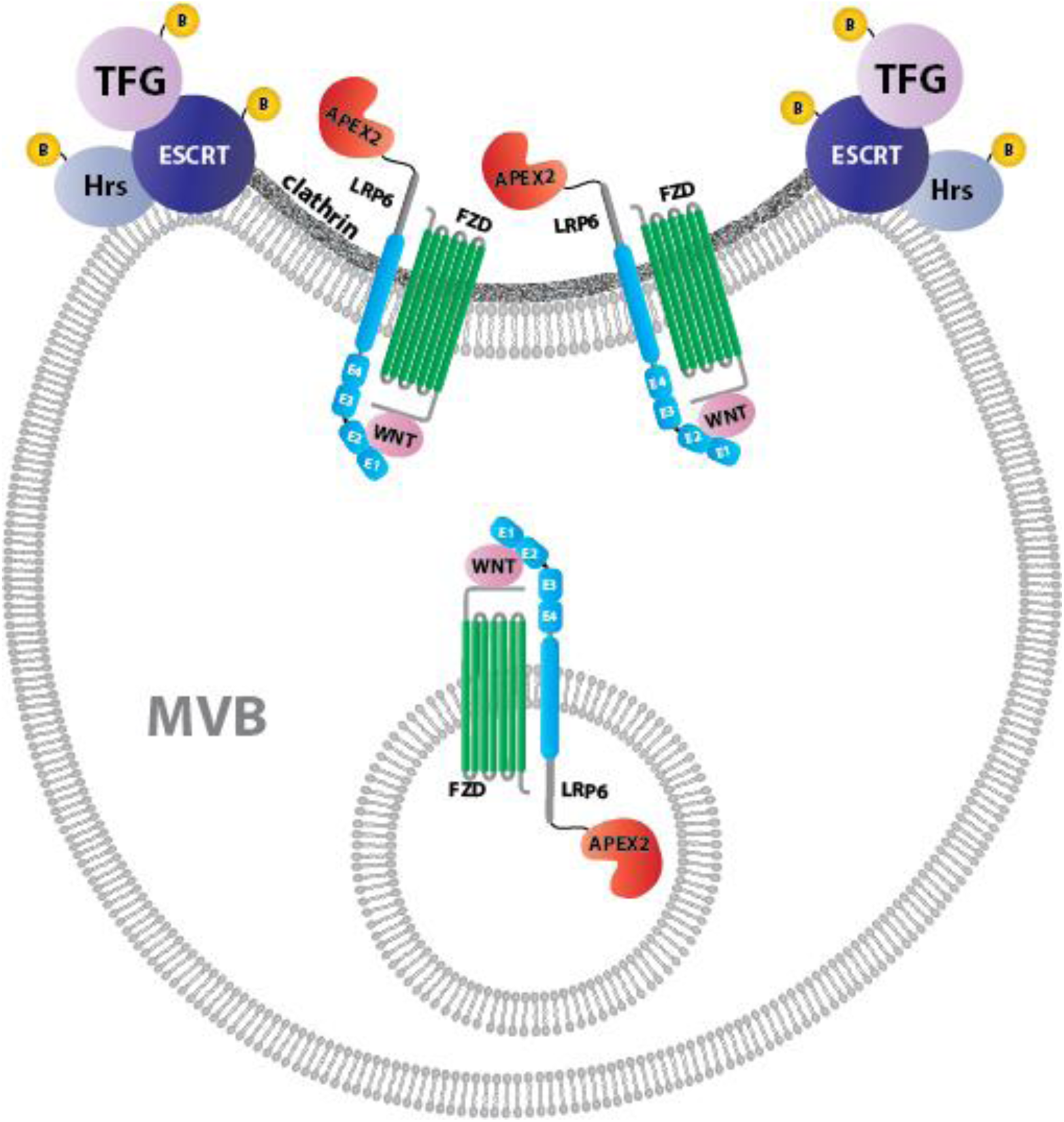
Model of how Lrp6-APEX2 may interact with the ESCRT machinery and TFG in multivesicular body formation during Wnt-induced endocytosis. The endocytosed Wnt receptor complex on the surface of multivesicular endosomes becomes clustered in clathrin-containing invaginations that give rise to the intraluminal vesicles (ILVs) of late endosomes. MVBs are the obligatory pathway through which plasma membrane proteins reach the lysosome. We found that most ESCRT GO components are brought into proximity of Lrp6-APEX2 5 min after treatment with Wnt3a conditioned medium. The top interactor with Lrp6-APEX2 after 20 minutes of Wnt3a was TFG, which we show here is required for Wnt signaling. TFG protein co-localizes with the ESCRT-0 protein Hrs/Hgs after Wnt stimulation. We did not detect interactions with intraluminal vesicle components such as CD63; a possible explanation is that the acidic medium or lysosomal proteases might interfere with APEX2 activity.

The connection between Wnt and endocytosis is well known (48, 49), and several factors have been found to modulate Wnt through endosomal mechanisms. For example, the protein arginine methyltransferase 1 (PRMT1), which interacts with Lrp6 (50), was shown to be indispensable for the sequestration of GSK3 into MVBs by microautophagy (15, 35) in order to sustain canonical Wnt signaling. In recent work, a proteomic proximity approach similar to the one reported here for Lrp6-APEX2, made use of a Fzd9b-APEX2 fusion to discover that Epidermal Growth Factor Receptor (EGFR)-mediated phosphorylation of Fzd9b was required for receptor internalization and signal transduction in response to Wnt9a (51). In that study, the proteomic data showed that endosome-associated Rabs and the Clathrin-endocytic pathway interacted with Fzd9b upon Wnt9a signaling, supporting the view that the endosomal machinery is an early component of the Wnt/β-catenin pathway. Importantly, in this study we found that Kras, which is a major regulator of macropinocytosis (52), interacts with Lrp6-APEX2 after Wnt treatment. A central role for endocytosis in Wnt signaling is consistent with the observations reported here, which revealed striking interactions with the ESCRT machinery that generates MVBs after Wnt treatment (Fig. 2B). In addition, Wnt3a treatment induced interactions with the actin cytoskeletal machinery, which is essential for Wnt-induced macropinocytosis (16, 37).

### Proximity Labeling with Lrp6-APEX2 Identifies a Novel Component of the Wnt Pathway

The best validation of the proximity biotinylation approach was the identification of an unexpected player in the Wnt pathway. Among the most enriched Lrp6 interactors we found TFG, a protein that was first identified in papillary thyroid carcinomas as a result of an in-frame oncogenic rearrangement between its N-terminal region and the tyrosine kinase domain of Neurotrophic tyrosine kinase receptor type 1 (NTRK1, also designated as Trk) (38). The ability to self-oligomerize, conferred by the TFG N-terminal coiled-coil domain, leads to constitutive activation of the tyrosine kinase domain responsible for the transforming activity of this chimeric oncogenic protein (39). Similarly, translocations between the TFG and anaplastic lymphoma kinase (ALK) genes result in a fusion oncoprotein in some lung cancers (53). The physiological function of wild type TFG has been previously investigated. Studies in *C*. *elegans* and human cultured cells reported that TFG localizes to the ER exit sites adjacent to the ER-Golgi intermediate compartment known as the ERGIC compartment, a cluster of membranes that secretory cargos traverse on their route to the Golgi apparatus (54). TFG hexamers were shown to co-localize and interact with the coat protein complex II (COPII) components Sec16 and Sec23, promoting the trafficking and secretion of cargo proteins, including synaptobrevin (54, 55). The importance of TFG in membrane traffic is also highlighted by the presence of several pathogenic mutations that impair TFG activity in ER function and are linked to different neurological disorders, including a hereditary motor and sensory neuropathy (HMSN-P) (56).

Our findings revealed an additional function for TFG, a protein that localized to the same endosomal vesicles as Hrs. Wnt3a treatment, which increases the number of late endosomes/MVBs (15, 16), stimulated the co-localization of TFG and HRS. Interactions between TFG, Hrs and other components of the ESCRT machinery such as Tsg101 have been reported on BioGRID (https://thebiogrid.org/), a repository for protein-protein interactions that curates data derived from high-throughput datasets. Moreover, β-catenin reporter assays in cell culture showed that TFG is required for Wnt signaling, as confirmed both by siRNA mediated knock-down and CRISPR/Cas9 knock-out of TFG. Our Cas9 genome editing experiments generated a cell line expressing only the first 90 amino acids of TFG. The resulting short protein lacks the coiled-coil domain, which spans from amino acids 97 to 124 (38). This detail is of importance, as the coiled coil domain is involved in TFG oligomerization and is required for its function and transforming activity (39), suggesting that our truncated TFG is a null mutant that resulted in a 50% inhibiton of canonical Wnt signaling.

The requirement for TFG during Wnt signaling is strongly supported by experiments in *Xenopus laevis* embryos. *In vivo* loss of function experiments using a TFG morpholino showed that TFG was required for correct antero-posterior (A-P) patterning and head development in frog embryos. In *Xenopus*, A-P patterning is regulated by an activity gradient of Wnt signaling which is lower in the rostral (head) region of the embryo (57). Numerous secreted Wnt inhibitors, including Dkk1 (30), Frzb (58) and Bighead (31) are required to maintain low levels of Wnt and allow head development. Embryos injected with TFG morpholino developed larger heads and cement glands, mimicking the effects of Dkk1 overexpression and pointing to a reduction of the endogenous Wnt signal. In agreement with this view, TFG knock-down also abolished the posteriorizing activity exerted by Wnt8 DNA overexpression or Bighead knockdown, showing *in vivo* that Wnt signaling requires TFG. At present, the molecular mechanism by which TFG regulates the Wnt pathway is unknown, but the co-localization with the ESCRT-0 component Hrs points to a role in the MVB compartment (Fig. 6). Future work will determine whether other Wnt-induced interactions of Lrp6, provided here as an open resource to the community in Dataset S1 Tab 1, also participate in canonical Wnt signaling.

## Materials and Methods

### Cloning and plasmids

pCS2 was used as a backbone for cloning recombinant DNA. First, the basic pCS2 plasmid was made compatible with the Gateway™ recombination system, by cloning a ccdb/chloramphenicol cassette (Invitrogen™, 11828029) into the pCS2 StuI restriction site. Flag-tagged cytosolic APEX2 (Addgene, 49386) was then amplified by polymerase chain reaction (PCR) and sub-cloned in frame with the 3’ of the Gateway™ cassette, into the pCS2 XhoI/XbaI sites. VSV-G-tagged human Lrp6 was obtained from Addgene (27282), amplified by PCR using primers containing Gateway sequences, and sub-cloned through a Gateway reaction following manufacturer’s instructions. Finally, an IRES sequence followed by a GFP-tagged Puromycin N-Acetyltransferase (PAC) gene, obtained from Addgene plasmid 45567, was amplified by PCR and sub-cloned into XbaI/SnaBI sites, downstream of the Lrp6-APEX2 open reading frame (ORF), to allow for selection of stable transfectants with the antibiotic puromycin. A human TFG clone was obtained from Dharmacon™, sequence verified, and Gateway-cloned into a pCS2 vector containing a 3xFlag epitope tag at the C-terminus. *Xenopus laevis* TFG was PCR-amplified from cDNA obtained from frog eggs, Gateway-cloned into pCS2-3xFlag and pCS2-Myc-Streptact. All cloned sequences were verified by DNA Sanger sequencing (Genewiz^®^). Primer sequences are shown on Table S1. The plasmids used in this study will be made available on the Addgene plasmid repository. All animal experiments procedures were approved by the UCLA institutional Animal Research Committee.

### Stable line transfection

The pCS2 Lrp6-APEX2-IRES-GFP-PAC plasmid was transfected into Human Embryonic Kidney 293T (HEK293T) cells using Lipofectamine2000 (Thermo Fisher Sci.). 48 hours after transfection puromycin was added to the culture medium, at a final concentration of 4 ug/ml. Stable transfected cells were continuously expanded and selected with antibiotic for at least two weeks, after which monoclonal lines were derived by limiting dilution. Two independent clones were obtained, which maintained stable and uniform expression of the Lrp6-APEX2 and GFP-PAC chimeras after many cell passages.

### Mass spectrometry analyses

Affinity-purified biotinylated protein samples were reduced and alkylated using 5 mM Tris (2-carboxyethyl) phosphine and 10 mM iodoacetamide, respectively, and then proteolyzed by the sequential addition of trypsin and lys-C proteases at 37°C as described (33, 34). The digestion reactions were stopped by the addition of 5% formic acid, desalted using Pierce C18 tips (Thermo Fisher Sci.), and then dried and resuspended in 5% formic acid. Peptide digests were fractionated online using a 25 cm long, 75 µm inner diameter fused silica capillary packed in-house with bulk C18 reversed phase resin (1.9 µm, 100A pores, Dr. Maisch GmbH). A 140-minute water-acetonitrile gradient was delivered to a maximum of 80% buffer B using a Dionex Ultimate 3000 UHPLC system (Thermo Fisher Sci.) at a flow rate of 300 nl/minute (Buffer A: water with 3% DMSO and 0.1% formic acid and Buffer B: acetonitrile with 3% DMSO and 0.1% formic acid). Peptides were ionized using a distal 2.2 kV spray voltage and a capillary temperature of 275°C and analyzed by tandem mass spectrometry (MS/MS) using an Orbitrap Fusion Lumos mass spectrometer (Thermo Fisher Sci.). Data was acquired by a Data-Dependent Acquisition (DDA) method comprised of a full MS1 scan at 120,000 resolution followed by sequential MS2 scans (Resolution = 15,000) to utilize the remainder of the 3 second cycle time. Data analysis was performed using two discrete bioinformatic pipelines. The first analysis used the Integrated Proteomics pipeline 2 (Integrated Proteomics Applications, San Diego, CA) to generate peptide and protein lists that were quantified using spectra counting. In this case, MS2 spectra were searched using the ProLuCID algorithm against the EMBL Human reference proteome (UP000005640 9606) followed by filtering of peptide-to-spectrum matches (PSMs) by DTASelect using a decoy database-estimated false discovery rate of <1%. In the second approach, an in-house Galaxy-based pipeline was used to provide label-free MS1 scan intensity-based quantification for the samples. MS2 spectra were searched against the EMBL Human reference proteome (UP000005640 9606) using the MSGF+ search engine (59). False detection rates for evaluating the peptide-spectrum-match (PSMs) were determined using Percolator while protein identification confidence was estimated via the stand-alone implementation of FIDO such that analytes had respective score cut-off q-values at or below 0.01 at both PSM and protein level (60-62). The Integrated peak areas of extracted ion chromatograms across multiple runs were calculated for each peptide using the Skyline software platform (63). In order to calculate the fold change and determine the protein abundance changes, the MSStats R-package was used to normalize across runs using quantile normalization, summarize peptide-level intensities into a protein-level abundance, and perform statistical testing to compare protein abundance across conditions (64). The raw dataset generated by mass spectrometry has been deposited in the online mass spectrometry interactive virtual environment (MassIVE) repository resource, with accession number MSV000084335. Additional methods, including proximity labeling, embryo and cell culture procedures are available in *SI Appendix, Supplemental SI Materials and Methods*.

## ACKNOWLEDGEMENTS

We thank Dr. Randall Moon, Dr. Christof Niehrs and Dr. Sylvie Urbé for reagents used in this study. We thank members of the E.M.D.R. laboratory for comments on the manuscript. L.V.A. was supported by an NIH postdoctoral fellowship (NIH F32 GM123622), N.T-M. by a UC MEXUS postdoctoral fellowship FE-17-65, and Y.J-A and J.A.W. by NIH GM089778 and GM112763. This work was also supported by the Norman Sprague Endowment and the Howard Hughes Medical Institute.

## Supporting Information for

### This PDF file includes

Figures S1-S5 and Legends

Tables S1, S2, S3

Captions for Movie S1

Supplemental SI Materials and Methods

Supplemental References

### Other supplementary materials for this manuscript include the following

Movie S1

Dataset S1

**Fig. S1.**
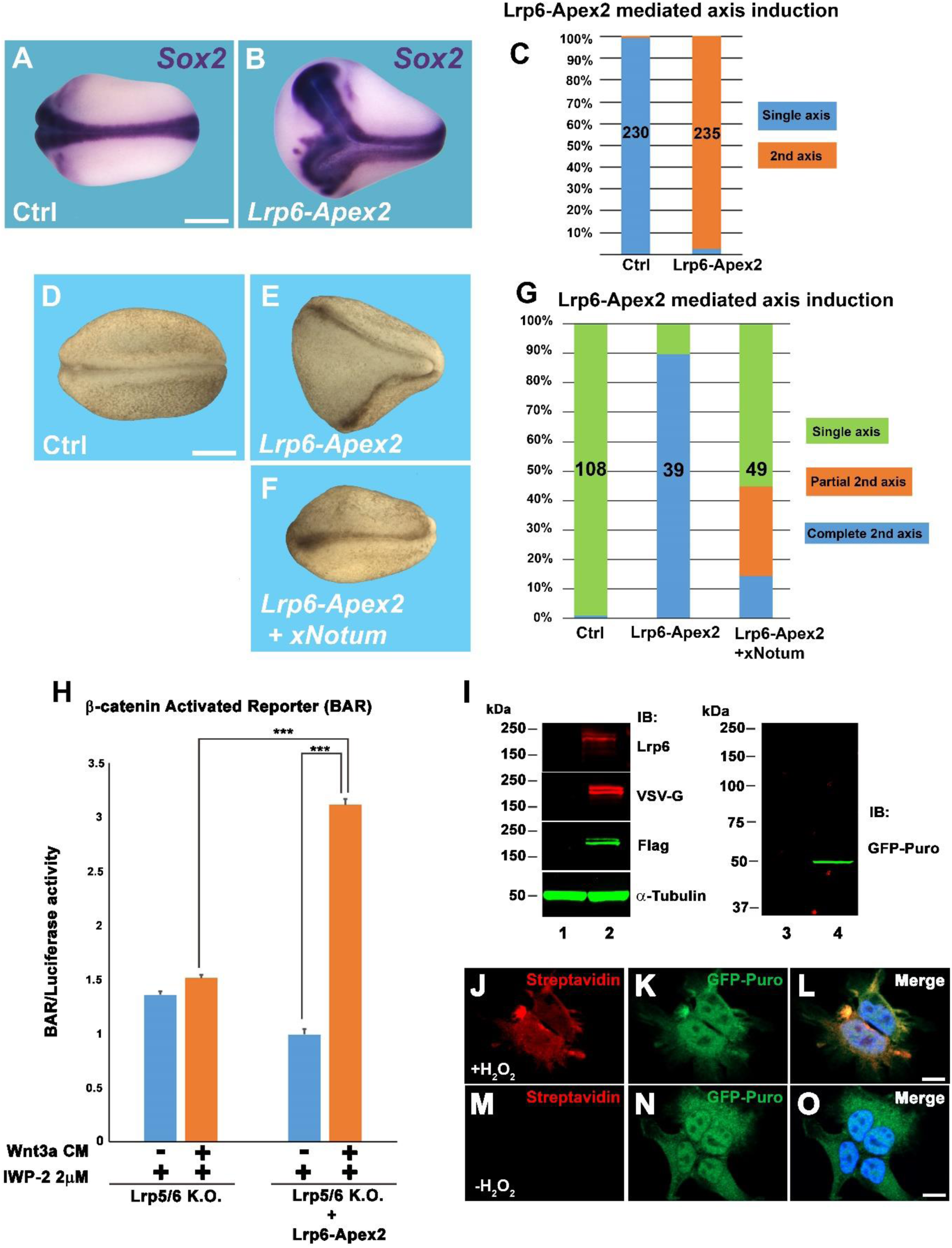
*Xenopus* embryo and cell-based assays demonstrate Lrp6-APEX2 transduces Wnt signaling. (*A, B*) Microinjection of 1 ng of *Lrp6-APEX2* mRNA into one ventral blastomere of *Xenopus* embryos at the 4-cell stage induced formation of a secondary axis, visualized through *in situ* hybridization for the neural marker *Sox2* at stage 20. Scale bar corresponds to 500 µm. (*C*) Quantification of the experiment shown in panels *A* and *B*. Lrp6-APEX2 induced double axes with a penetrance of almost 100%. The number of embryos used for these experiments are shown inside the columns. (*D*-*F*) Injection of 1 ng of *Lrp6-APEX2* mRNA into a single ventral blastomere of *Xenopus* embryos at the 4-cell stage induced secondary axes, but this required endogenous Wnt ligands as co-injection of 300 pg of mRNA encoding the Wnt de-acylase *Notum* strongly inhibited secondary axis formation. Embryos were fixed and scored at stage 20. Scale bar corresponds to 500 µm. (*G*) Quantification of the experiment shown in panels *D*-*F*. Lrp6-APEX2 induced double axes with a penetrance of 90%, and co-injection with the Wnt inhibitor *Notum* strongly inhibited this activity. (*H*) β-catenin-Activated reporter (BAR)-Luciferase assay showing that transient expression of Lrp6-APEX2 was able to rescue response to Wnt3a conditioned medium in Lrp5/6 double knock-out cells (1). Double knock-out cells were preincubated overnight with 2 µM (final concentration) of the Porcupine inhibitor IWP-2 to prevent activation by endogenous Wnt signaling in the rescued cells. (*I*) Western Blot of whole cell lysates derived from parental HEK293T cells (lanes 1 and 3) or Lrp6-APEX2 293T clonal cells (lanes 2 and 4). Robust Lrp6 expression was detected with different anti-Lrp6 antibodies (Lrp6, VSV-G and Flag) in this permanent cell line. Note the presence of the two typical bands, with the lower one corresponding to the immature form of Lrp6, α-tubulin was used as a loading control. Lane 4 shows correct expression of a GFP-tagged selectable marker, puromycin N-acetyl-transferase, whose expression is driven by an IRES placed downstream of Lrp6-APEX2. (*J*-*O*) APEX2 peroxidase activity in Lrp6-APEX2 293T cells could be visualized by fluorescent Streptavidin, which binds to proteins biotinylated by APEX2. Note that Streptavidin fluorescence was observed only in presence of H_2_O_2_ (and biotin-phenol), which is the substrate for peroxidase activity. GFP-puromycin was used as counter-staining, and DAPI nuclear staining is shown in the Merge panels. Scale bars represent 10 µm.

**Fig. S2.**
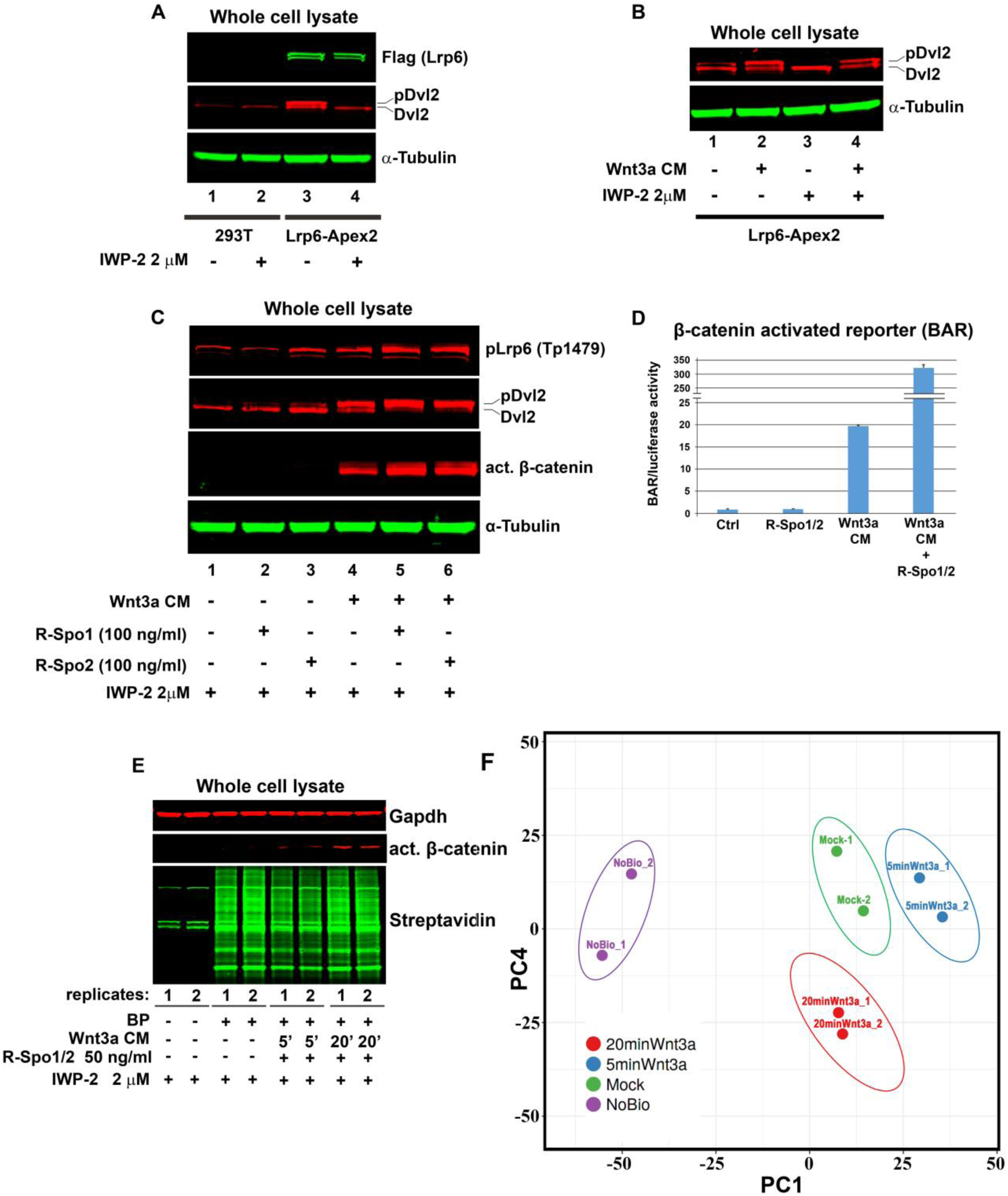
Lrp6-APEX2 cells respond to exogenous Wnt signals in the presence of IWP-2, and biotinylation of key Wnt pathway proteins after Wnt addition. (*A*) Western blot of lysates from control HEK293T and Lrp6-APEX2 cells. Lrp6-APEX2 cells show high levels of phosphorylated Dvl2 (pDvl2), a hallmark of activated Wnt signaling (lane 3), even in absence of exogenous Wnt. However, overnight incubation with 2 µM IWP-2, an inhibitor of the Wnt-palmitoyltransferase Porcupine, prevented Dvl2 phosphorylation (compare lanes 3 and 4). pDvl2 was not detected in 293T control cells (lanes 1 and 2) even in the absence of IWP-2 incubation, indicating that permanent transfection with Lrp6-APEX2 sensitizes cells to low levels of endogenous Wnt signals; α-tubulin was used as a loading control. (*B*) Western blot of Lrp6-APEX2 293T cell lysates. In absence of IWP-2, pDvl2 was detected regardless of Wnt3a conditioned medium (CM) treatments (lanes 1 and 2). However, Lrp6-APEX2 cells incubated overnight with IWP-2 showed Dvl2 phosphorylation only in the presence of Wnt3a CM (lanes 3 and 4); α-tubulin was used as a loading control. (*C*) Western blot of Lrp6-APEX2 cell lysates showing that cells responded robustly to Wnt3a CM and that this was further potentiated by R-spondins 1 and 2, even in the presence of Porcupine inhibitor IWP-2. Wnt3a CM induced robust Dvl2 phosphorylation, stabilization of non-phosphorylated (active) β-catenin, and Lrp6 phosphorylation at Threonine residue 1479 (compare lanes 1 and 4) after 3 hours of Wnt3a treatment. Wnt pathway activation was further increased when R-Spondin 1 and 2 recombinant proteins (2) were added to Wnt3a CM (lanes 5 and 6). R-Spo proteins alone had little effect on Wnt activation (because of the presence of IWP-2); α-tubulin was used as a loading control. (*D*) Wnt3a CM activity was tested on a permanently transfected HEK293T β-catenin activated reporter (BAR) cell line. Wnt3a CM induced a 20-fold increase in luciferase activity, while combination of Wnt3a CM and R-Spo1/2 proteins (50 ng/ml each) induced a 200-fold increase. This was adopted as the fortified Wnt3a CM formulation was used for proteomic experiments. (*E*) Proximity biotin-labelling of Lrp6-APEX2 cell in aliquots of the same large-scale cultures used in the proteomic experiments after 5 min or 20 min of treatment with Wnt3a CM containing recombinant R-Spo1/2 (50 ng/ml each). Western blot using Streptavidin-IRDye 800 was performed to confirm that biotinylation occurred (note that the biotin “ladder” was observed only in presence of Biotin-phenol, confirming specificity). The three bands seen in lanes 1 and 2 represent endogenous biotin-containing carboxylases. Non-phosphorylated Ser33/37/Thr41 β-catenin, which represents its active form (act. β-catenin) (3-5), confirmed Wnt pathway activation; Gapdh was used as a loading control. Experiments were conducted on biological duplicates, which were then processed for mass spectrometry. (*F*) Principal Component Analysis (PCA) to analyze the dimensionality and trends exhibited by the enrichment profiles of the different proteomic experiments. The samples included biological duplicates of 4 different cell treatments: no-biotinylation control (NoBio), treatment with control CM (Mock) and treatment with Wnt3a CM for 5 minutes (5minWnt3a) or 20 minutes (20minWnt3a). Each axis represents a principal component (PC1 and PC4). These PCA analyses confirmed reproducibility of experiments by clustering duplicate conditions. Each dot represents a sample and each color represents the type of treatment.

**Fig. S3.**
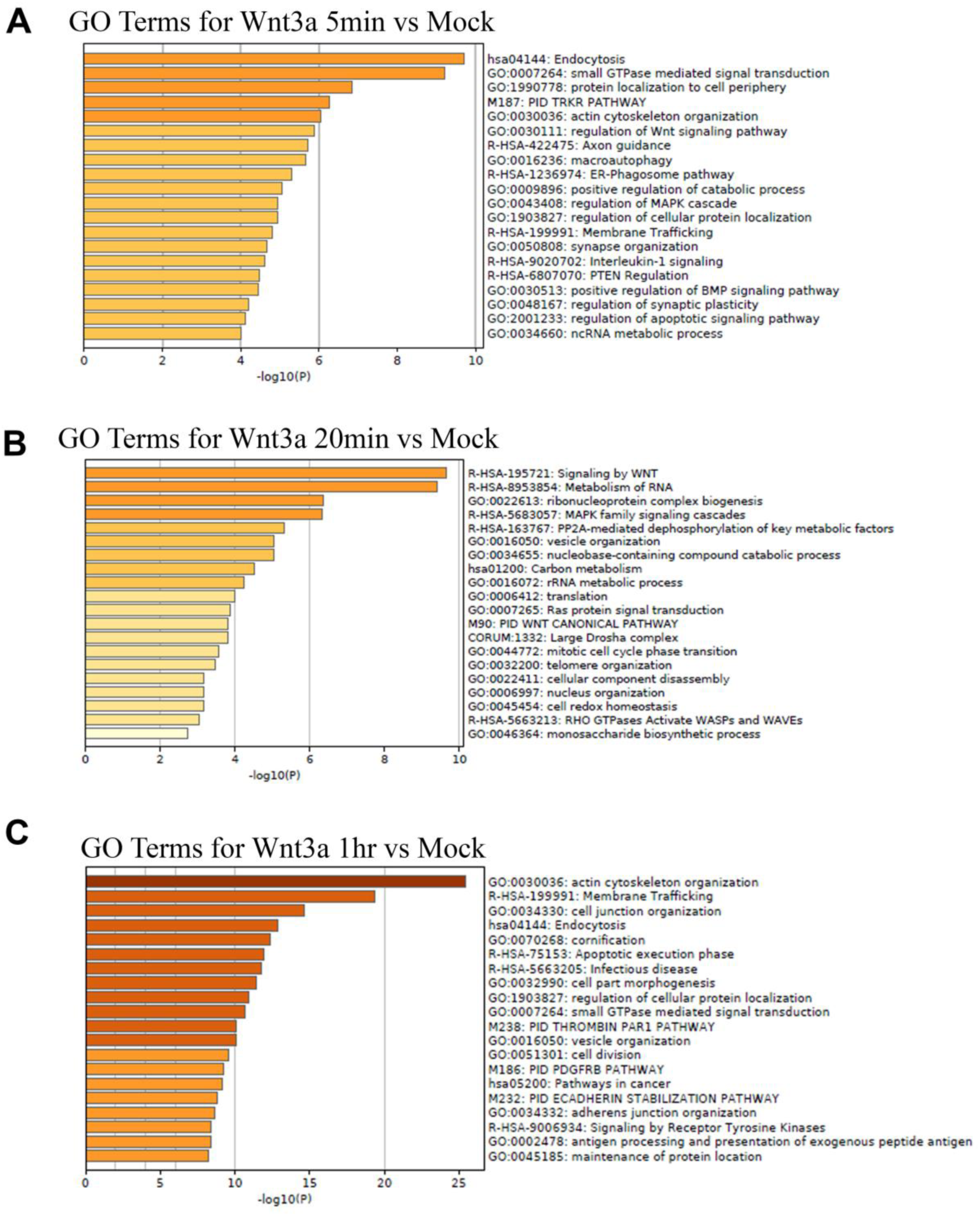
Gene Ontology analyses of biotinylated Wnt-induced Lrp6-APEX2 target proteins at three timepoints. Biotinylated proteins from Wnt samples were ranked for enrichment against the control Mock samples (cells treated with control medium). The top hits (150-300 top proteins) were then analyzed with the Metascape online software (6), available at metascape.org/gp/index.html#/main/step1. (A) Analysis of the top 150 biotinylated Lrp6 interactors after 5 minutes of Wnt3a treatment. (B) Analysis of 130 biotinylated Lrp6 interactors after 20 minutes of Wnt3a treatment. (C) Analysis of 300 top biotinylated Lrp6 interactors after 1 hour of Wnt3a treatment. Bars represent the enriched terms across input protein lists, colored by p-values.

**Fig. S4.**
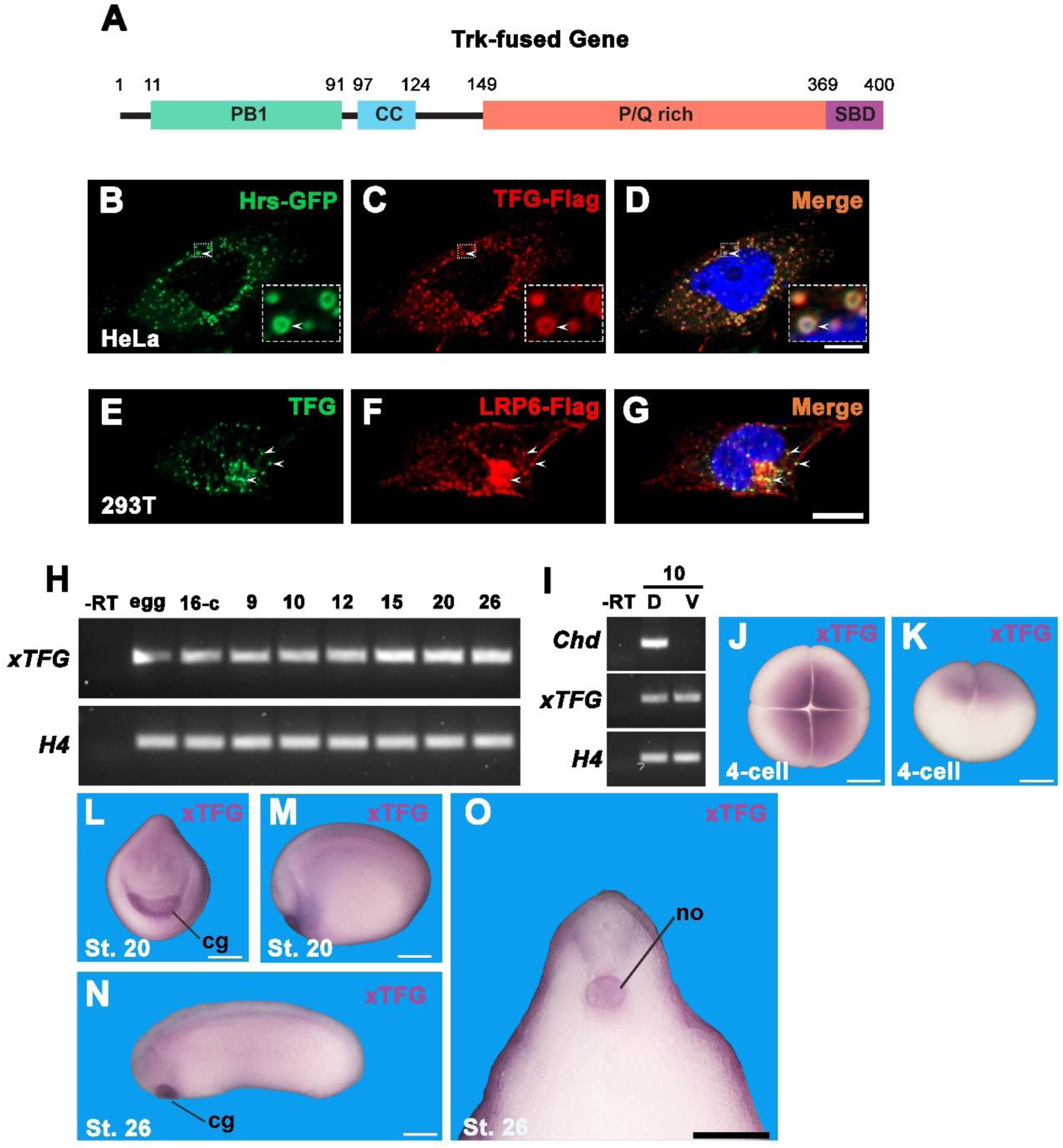
TFG colocalizes with Hrs and Lrp6 and is expressed in developing *Xenopus laevis* embryos. (*A*) Diagram highlighting human TFG protein structure and its structural domains (7). PB1, also called Phox and Bem1p domain, is possibly involved in heterodimer formation and is required for oncogenic activity of TFG fusion products; CC is a coiled-coil domain involved in oligomerization and also required for the transforming activity of oncogenic TFG fusions; P/Q rich is a Proline/Glutamine rich domain, a disordered region required for localization of TFG at the endoplasmic reticulum (ER)-Golgi intermediate compartment (ERGIC). SBD is the Sec23-binding domain, required for interaction with Sec23 and disassembly of COPII vesicles. Numbers indicate amino acid position. (*B*-*D*) Immunofluorescence on HeLa cells transfected with plasmid DNAs encoding mouse Hrs-GFP and human TFG-Flag. Note the strong colocalization between the two proteins. Inset shows overlay at the periphery of endocytic vesicles. These results strongly support the specificity of endogenous HRS and TFG immunostainings shown in Fig. 3 of the main text. DAPI nuclear staining (in blue) is shown in Merge panel; scale bar represents 10 µm. (*E*-*G*) Immunofluorescence of Lrp6-APEX2 293T cells showing partial endogenous partial overlap with endogenous TFG (arrowheads), indicating co-localization between the two proteins and corroborating the results from proteomic analysis. DAPI nuclear staining (in blue) is shown in Merge panel; scale bar represents 10 µm. (*H*) RT-PCR analysis on cDNA obtained from *Xenopus* embryos at different developmental stages. *xTFG* mRNA was present as a maternal factor (in eggs and 16-cell stage embryos) and continued to be expressed at later stages. *Histone 4* (*H4*) was used as a loading control. -RT was used as a negative control. (*I*) RT-PCR on dorsal (D) and ventral (V) fragments dissected from stage 10 *Xenopus* gastrulae. Note that *xTFG* transcripts do not show any dorso-ventral preference, unlike the organizer gene C*hordin* (*Chd*). *Histone 4* (*H4*) was used as a loading control and -RT as a negative control. (*J, K*) Whole mount *in situ* hybridization showing animal pole localization of *xTFG* maternal transcripts, at the 4-cell stage. *J* offers a top-view and *K* a side-view of the same embryo. (*L*-*N*) *xTFG* mRNA was expressed strongly in the cement gland (cg) at later stages of development (stage 20 and 26). Weaker staining could also be detected in the epidermis. (*O*) Bisected stage 26 embryos revealed specific *xTFG* mRNA staining in the notochord (no). Weaker expression was detected in the epidermis and the alar plate of the spinal cord. Scale bar represents 500 µm in *J*-*N*, and 200 µm in *O*.

**Fig. S5.**
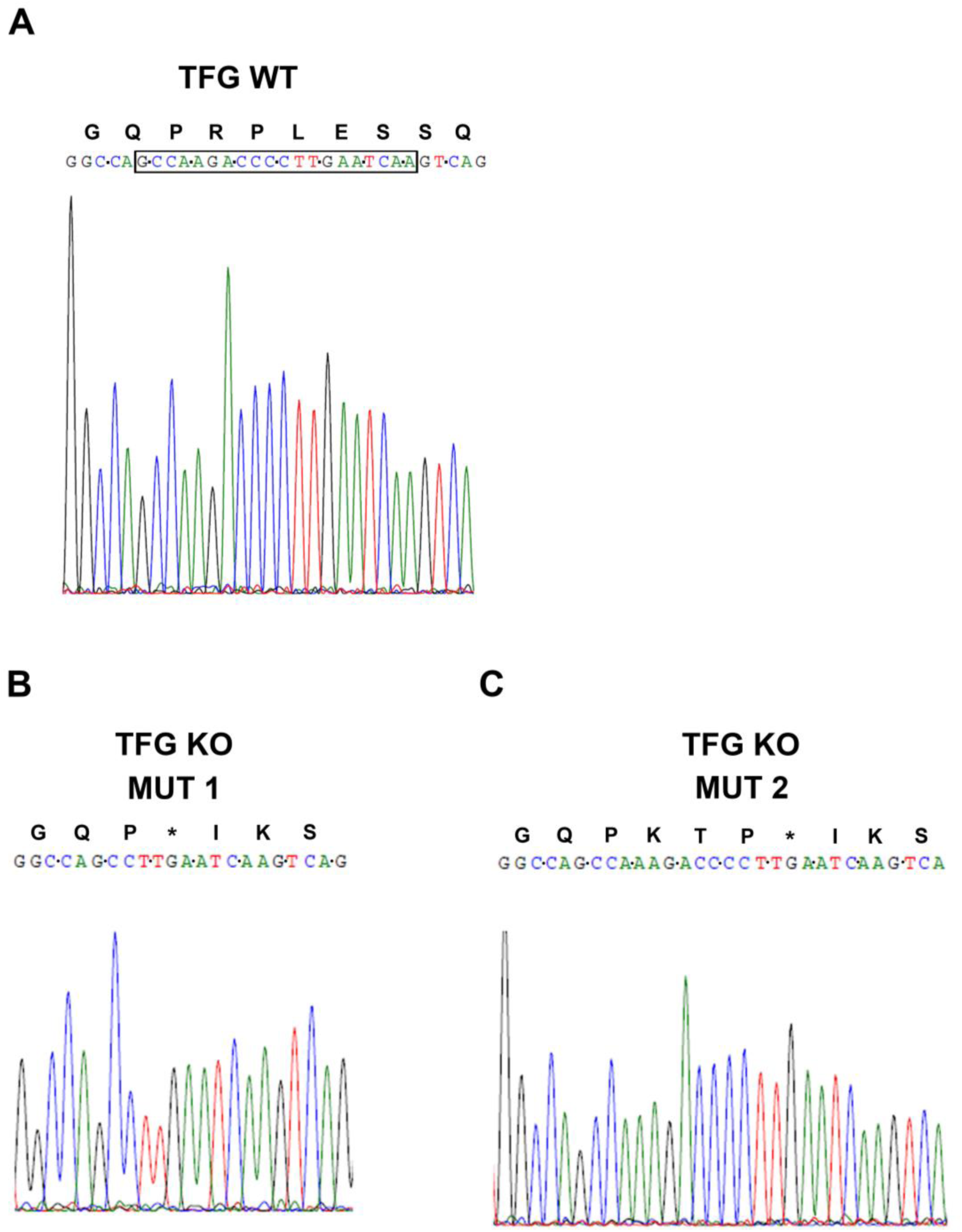
TFG knock-out by CRISPR-Cas9 confirmed by genomic DNA sequencing. Genomic DNA was extracted from untransfected WT HEK293T cells, and from a cell clone stably transfected with a Cas9 expression vector containing an sgRNA-encoding sequence specific for human TFG. A 600 nucleotide region spanning the target sequence was amplified by PCR, cloned into sequencing vectors, and analyzed by Sanger sequencing. The reference chromatogram from sequencing results is shown below the DNA sequence. (*A*) Nucleotide sequence of TFG from wild-type (WT) HEK293 cells; the coding frame is indicated by black dots between each codon. The boxed area marks the target sequence recognized by the spacer region in the sgRNA in the WT TFG sequence. The predicted amino acid sequence is shown above the DNA sequence. (*B* and *C*) Sequences from a TFG knock-out clonal cell line showing the two types of mutant sequences resulting after PCR amplification of mutated genomic DNA. Each TFG allele harbored independent insertion-deletion (indel) events induced by Cas9 as a result of non-homologous end joining (NHEJ) repair. In Mutation 1 (Mut1) Cas9 induced a deletion of a short nucleotide sequence (8 nucleotides, AAGACCCC). Mutation 2 (Mut 2) resulted in the insertion of an extra adenine between the original CCA and AGA codons. Both indel events induced a frameshift mutation, introducing an early STOP codon indicated by an asterisk, generating truncated TFG mutant proteins of approximately 90 amino acids.

**Fig. S6.**
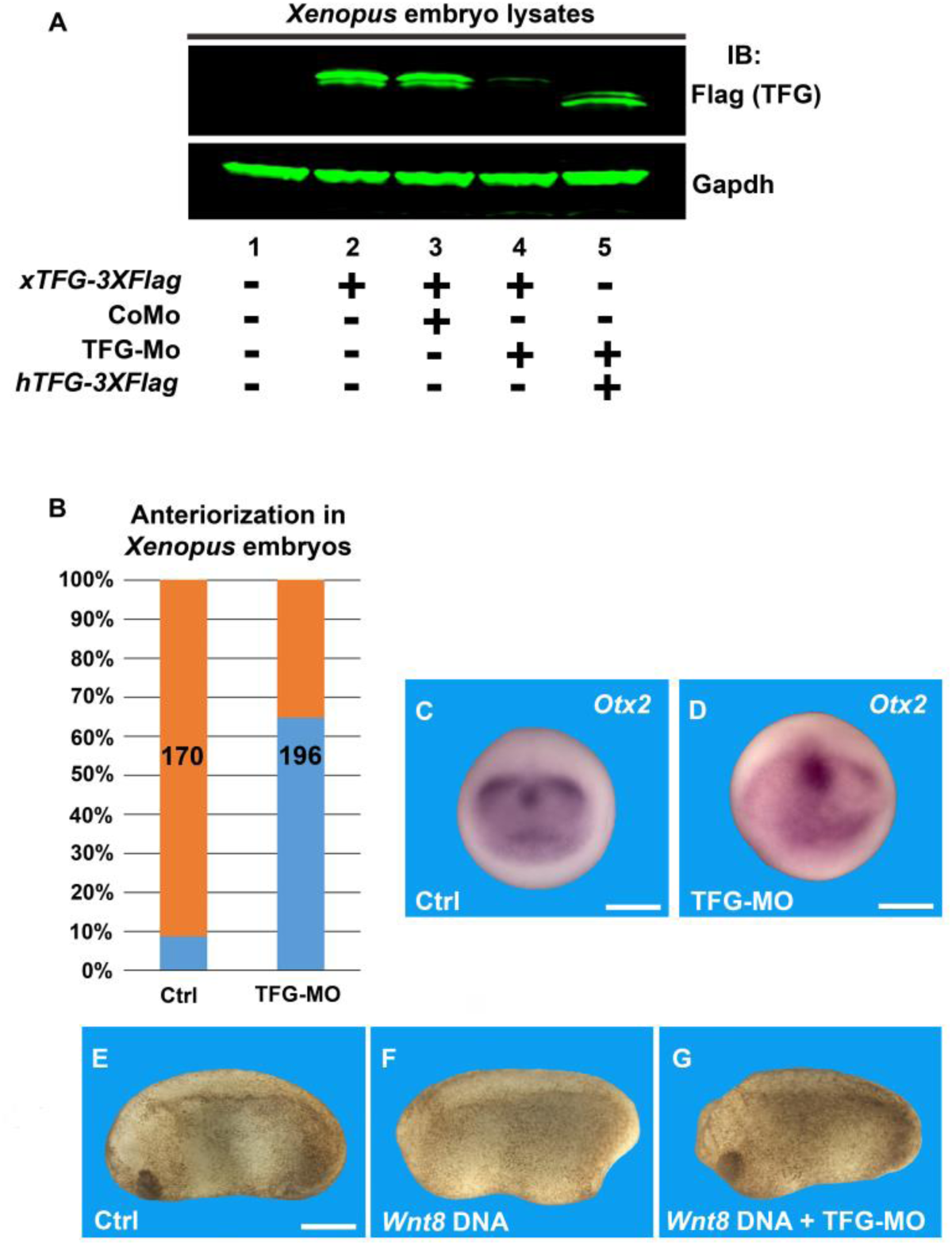
TFG loss-of-function using a specific antisense Morpholino (MO) induces anteriorization, revealing a requirement for Wnt activity in *Xenopus* embryos. (*A*) TFG-MO is specific and blocked translation of Flag-tagged *Xenopus TFG* mRNA in microinjected embryos (compare lanes 3 and 4), while mRNA encoding human *TFG* was not targeted by the morpholino (lane 5). Frog embryos were injected at 4-cell as indicated, cultured until gastrula stage and lysed for Western blot analysis. 400 pg/embryo of *Xenopus* or human Flag-tagged *TFG* mRNA were microinjected together with 72 ng/embryo of Control Morpholino (CoMO) or xTFG-MO. Uninjected embryos were used as a negative control (lane 1), and Gapdh used as loading control. (*B*) Quantification of the experiment shown in Fig. 5 *B* and *C*. Numbers in the columns show the amount of embryos used. TFG-MO induces anteriorization in over 60% of the injected embryos. (*C, D*) *In situ* hybridization for the anterior marker *Otx2*. Compared to control embryos (n=33), TFG-MO induced an expansion of *Otx2* in 65% of the injected embryos (n=20). (*E*-*G*) While control embryos showed normal antero-posterior patterning (n=103), but injection of *Wnt8* DNA induced loss of anterior development, as shown by the absence of the cement gland (90%, n=61). Co-injection of TFG-MO inhibited anterior truncation (65%, n=66), suggesting that Wnt signaling requires TFG. Scale bars in *C*-*D* represent 500 µm.

**Table S1.**
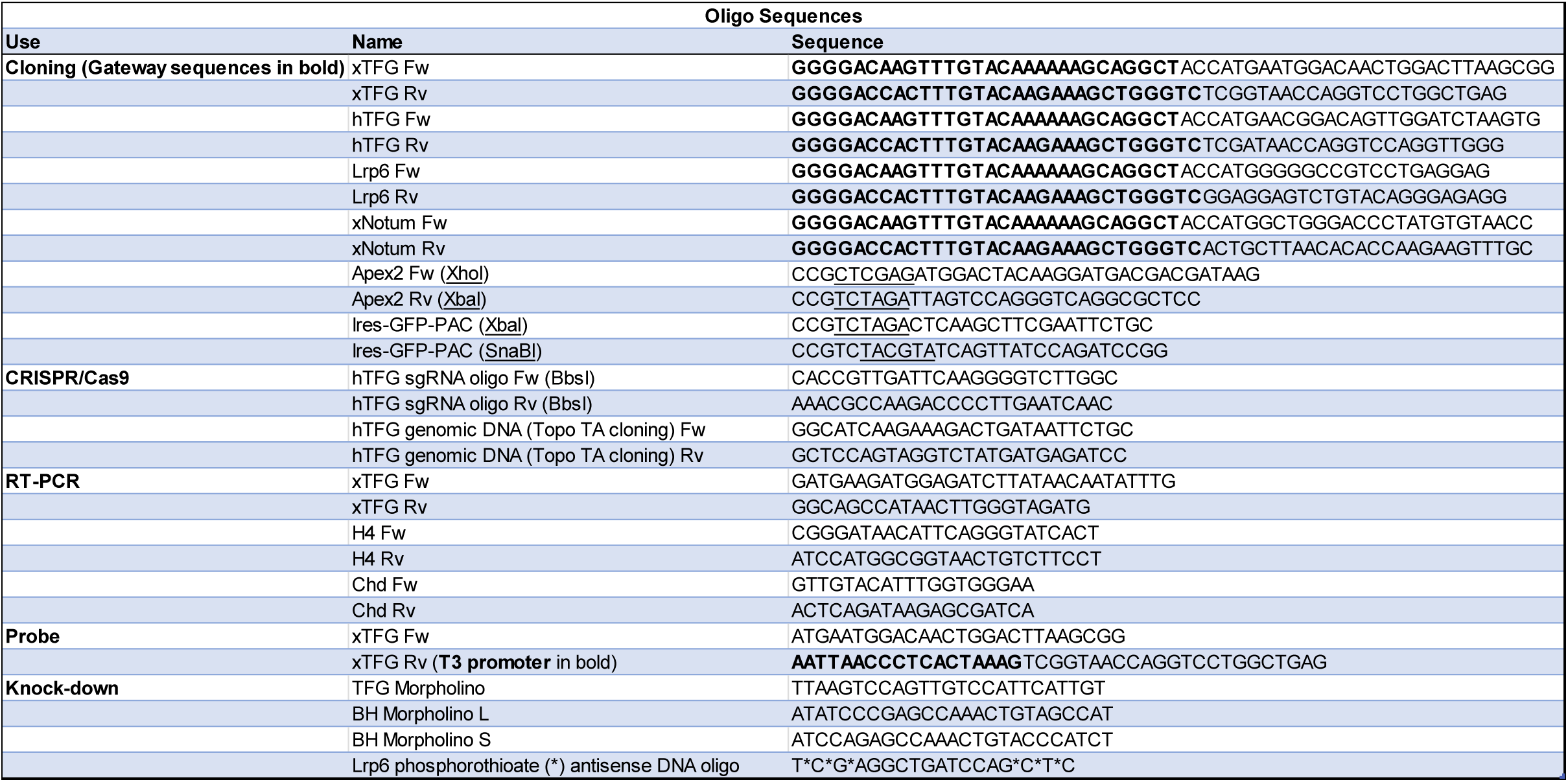
List of oligonucleotides used in this study.

**Table S2.** List of primary and secondary antibodies used for Western Blot in this study.

| **Western Blot** |  |  |  |
| --- | --- | --- | --- |
| **Primary Antibody** | **Vendor** | **Catalog Number** | **Dilution** |
| Rabbit anti-VSV-G | Genetex | GTX18612 | 1:1000 |
| Mouse M2 anti-Flag | Sigma | F3165 | 1:1000 |
| Rabbit anti-Lrp6 | Cell Signaling Technology | C5C7 | 1:1000 |
| Rabbit anti-phospho-Threonine 1479 Lrp6 | Abnova | PAB12632 | 1:1000 |
| Rabbit anti-TFG | Invitrogen | MA5-32375 | 1:1000 |
| Mouse anti-α-tubulin | Calbiochem | CP06 | 1:3000 |
| Chicken anti-GFP | Aves Labs | 1020 | 1:1000 |
| Rabbit anti-Gapdh | Cell Signaling Technology | 14C10 | 1:3000 |
| Rabbit anti-Dvl2 | Cell Signaling Technology | 30D2 | 1:1000 |
| Rabbit anti-active (non-phosphorylated) β-Catenin | Cell Signaling Technology | D13A1 | 1:1000 |
| Rabbit anti-β Actin | Cell Signaling Technology | 13E5 | 1:3000 |
| Streptavidin IRDye 800 CW | Li-COR | 926-32230 | 1:3000 |
| **Secondary Antibody** |  |  |  |
| Donkey anti-rabbit IRDye 680 RD | Li-COR | 926-68073 | 1:3000 |
| Donkey anti-mouse IRDye 800 CW | Li-COR | 926-32212 | 1:3000 |
| Donkey anti-chicken IRDye 680 RD | Li-COR | 925-68075 | 1:3000 |

**Table S3.** List of primary and secondary antibodies used for Immunofluorescence in this study.

| Immunofluorescence |  |  |  |
| --- | --- | --- | --- |
| Primary Antibody | Vendor | Catalog Number | Dilution |
| Mouse M2 anti-Flag | Sigma | F3165 | 1:300 |
| Mouse anti-TFG | Invitrogen | MA5-25759 | 1:200 |
| Mouse anti-β catenin | Santa cruz Biotech | sc-7963 | 1:200 |
| Chicken anti-GFP | Aves Labs | 1020 | 1:300 |
| Rabbit anti-Hrs | Santa Cruz Biotechnology | M-79 | 1:100 |
| Streptavidin Alexa Fluor 594 | Invitrogen | S11227 | 1:1000 |
| Phalloidin Alexa Fluor Plus 405 | Invitrogen | A30104 | 1:1000 |
| Secondary Antibody |  |  |  |
| Donkey anti-mouse Alexa Fluor 488 | Jackson ImmunoResearch Laboratories | 715-546-150 | 1:1000 |
| Donkey anti-rabbit Cy3 | Jackson ImmunoResearch Laboratories | 711-166-152 | 1:1000 |
| Goat anti-chicken Alexa Fluor 488 | Jackson ImmunoResearch Laboratories | 103-545-155 | 1:1000 |
| Donkey anti-mouse Cy3 | Jackson ImmunoResearch Laboratories | 715-166-150 | 1:1000 |

## *SI Appendix* Movie S1

**SI Appendix Movie S1.**
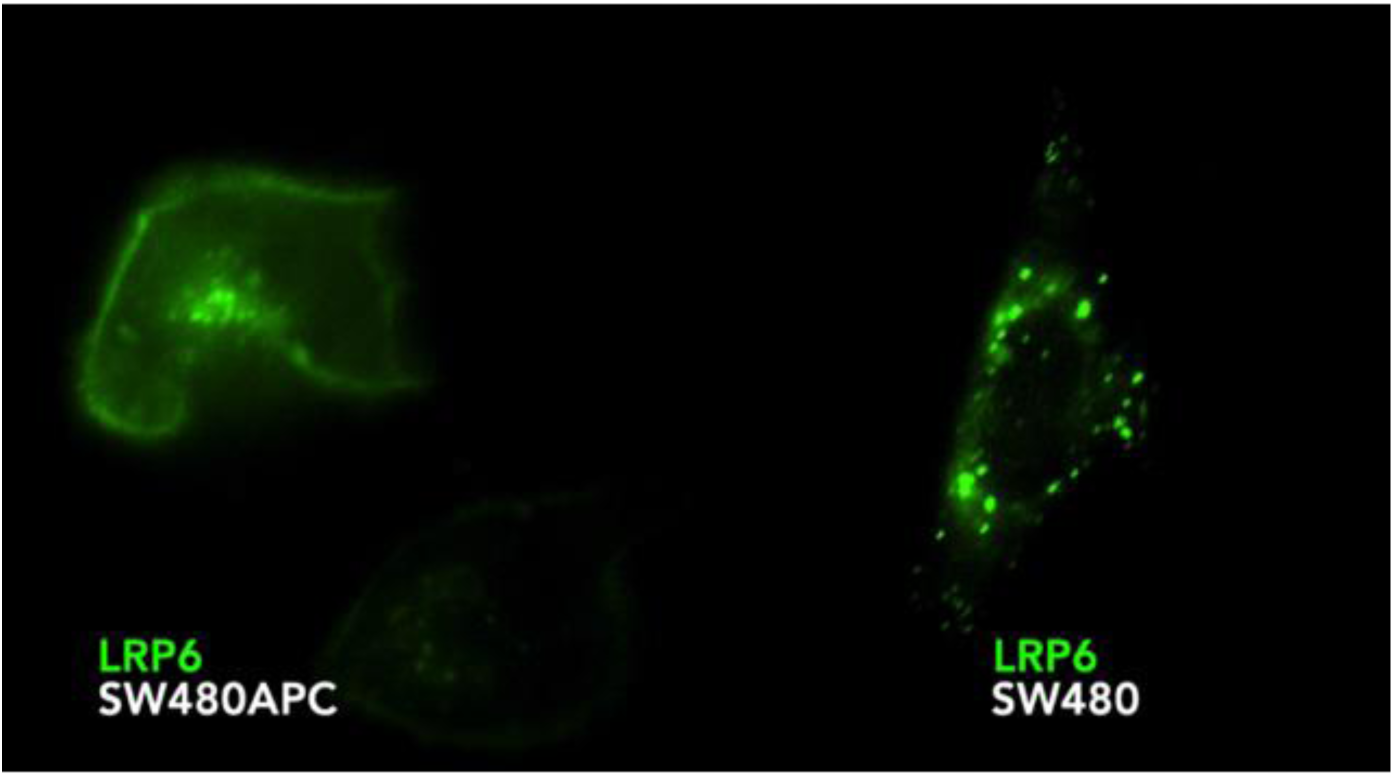
Lrp6 is constitutively endocytosed in SW480 colon cancer cells in which Wnt signaling is activated by mutation of Adenomatous Polyposis Coli, the tumor suppressor initially mutated in 85% of colon cancers, but not in cells in which APC is restored. SW480 cells were transfected with *pCS2-Lrp6-eGFP* DNA. Green fluorescence filter with Apotome.2 optical sectioning was used to collect images of Lrp6-eGFP. Note that in absence of APC (SW480 cell, right side) Lrp6-eGFP is found predominantly in intracellular puncta which represent endolysosomes. In SW480 cells in which APC has been restored Lrp6-eGFP localizes to the plasma membrane (SW480APC, left side). The only difference between these two cells is the presence or absence of APC. Wnt activation by loss-of-function of APC has profound effects on Lrp6 endocytic trafficking.

## Supplementary SI Materials and Methods

### APEX2-mediated proximity labeling

Lrp6-APEX2 HEK293T stable clonal cells cells were seeded in T225 flasks (Thermo Fisher Sci.) and cultured in filter-sterile growth medium (DMEM supplemented with 10% Fetal Bovine Serum (FBS), 1% penicillin/streptomycin and 1% glutamine), at 37 °C and 5% CO_2_. Importantly, the Porcupine inhibitor IWP-2 was also added to the medium, 2 µM final concentration, to prevent activation of Wnt signaling by ligands produced endogenously. When cells reached confluency, culture medium was replaced with 25 ml of growth medium supplemented with 500 µM Biotin-Phenol (BP, Iris Biotech). BP was omitted in the no-biotinylation (NoBio) control. For the Mock control samples, the medium was replaced again after 10 min with control conditioned medium (derived from control mouse L cells, ATCC^®^) containing 500 µM BP and cells were incubated for an additional 20 min. For the Wnt treatment samples, BP-containing medium was replaced after either 10 min or 25 min with Wnt3a conditioned medium (derived from mouse L Wnt3a cells, ATCC^®^) containing 500 µM BP, and cells incubated for an additional 20 min or 5 min, respectively. Thus, all samples were incubated with BP for a total of 30 min at 37°C. To start the biotin-labeling reaction, H_2_O_2_ was added directly to the BP-containing medium to a 1 mM final concentration 1 minute before the end of incubation and flasks were gently agitated. At the end of the 1-minute labeling reaction, cells were immediately washed 5 times with quencher solution (ice-cold Dulbecco’s phosphate-buffered saline, DPBS, containing 10 mM sodium azide, 10 mM sodium ascorbate and 5 mM Trolox). To minimize cell detachment during washes, flasks were gently inverted to pour the wash solution on the bottom face, and then inverted again to cover cells with fresh solution. Finally, cells were vigorously washed off the flasks, pelleted by brief centrifugation, and lysed in 1.5 ml of RIPA lysis buffer (50 mM Tris, 150 mM NaCl, 0.1% wt/vol SDS, 0.5% sodium deoxycholate, 1% vol/vol Triton X-100 in nanopure water, pH 7.5, supplemented with 1x Roche cOmplete™ protease inhibitor cocktail, 10 mM sodium azide, 10 mM sodium ascorbate and 5 mM Trolox). Two T225 flasks were used for each sample, and experiments were conducted in duplicates.

### Streptavidin pull-down

Cell lysates were cleared by centrifugation at 16,000 g for 10 min at 4°C. Supernatants were transferred to new 1.5 ml Eppendorf tubes and protein concentration of each sample was determined using the Pierce 660-nm protein assay, following manufacturer’s instructions. Streptavidin-coated magnetic beads (Pierce) were washed twice with RIPA buffer, and 2,000 µg of each protein lysate sample were then incubated with 100 µl of the magnetic bead slurry, on an end over end rotator for 1 hour at room temperature. Subsequently, the beads were washed twice with RIPA buffer, and an additional three times with RIPA buffer containing no Triton X-100/sodium deoxycholate since they interfere with subsequent mass spectrometry analysis, once with 1 M KCl, once with 0.1 M Na_2_CO_3_ pH 11, and three times with 8 M urea buffer in 100 mM Tris-HCl, pH 8.5. At this point, affinity-purified biotinylated protein samples were processed for further analysis by mass spectrometry.

### Generation of TFG knock-out cell lines by CRISPR-Cas9 genome editing

A plasmid based on the original PX459 from Zhang lab, containing Streptococcus pyogenes Cas9 (spCas9)-2A-blasticidin resistance was obtained from Addgene (#118055, from Ken-Ichi Takemaru lab). The human TFG genomic sequence was analyzed with the CRISPR Design tool available online at Synthego.com, to identify potential target sequences followed by the NGG protospacer adjacent motif (PAM) sequence. A promising 20-nt sequence on the reverse, non-coding strand was identified (5’-TTGATTCAAGGGGTCTTGGC<u>TGG</u>-3’, underlining indicates the PAM sequence) less than 300 nucleotides away from the TFG start codon. Oligos from both strands of the spacer TFG-specific sequence were annealed and phosphorylated following the directions from the Zhang lab protocol available at Addgene, https://www.addgene.org/crispr/zhang/. The double-stranded oligos were then cloned into spCas9-2A-blast, previously linearized with BbsI restriction enzyme (New England Biolabs). After transformation, plasmid DNA was extracted from bacterial colonies and analyzed by DNA sequencing for the presence of the sgRNA-encoding sequence. The resulting TFG-sgRNA spCas9-2A-blast plasmid was transfected with lipofectamine 2000 (Invitrogen) into HEK293T cells permanently transfected a Wnt-inducible β-catenin activated (BAR) Luciferase reporter and Renilla for normalization (see below). After 48 hours of transfection, permanently transfected cells were selected in medium containing 5 µg/ml blasticidin. Clonal lines were obtained by limiting dilution, expanded, and analyzed by western blot for TFG protein expression. One clone showed complete loss of TFG protein expression. To further confirm TFG gene KO, genomic DNA was extracted from mutant and WT cells and used as a template for PCR, using a proof-reading polymerase (Phusion high fidelity DNA polymerase, New England Biolabs). An amplification product of approximately 600 bp surrounding the Cas9 target sequence was obtained and ‘A-tailed’ (to add single A nucleotides at each 3’ end) using a standard non proof-reading taq polymerase (Taq 2X master mix, NEB). The tailed PCR product was then cloned into TOPO-TA cloning vector (Invitrogen), according to manufacturer instructions. Following transformation of the ligation reaction, multiple bacterial colonies were selected, grown, and plasmid DNA extracted and processed for sequencing.

### Embryo manipulations

*Xenopus laevis* frogs were purchased from Nasco. A sperm suspension was obtained after crushing 1 whole testis in 1 ml of 1x Marc’s modified ringers solution (MMR, 0.1 M NaCl, 2.0 mM KCl, 1 mM MgSO4, 2 mM CaCl2, 5 mM HEPES, pH 7.4). Eggs were spawn in high salt solution (1.2x MMR) from female frogs injected the night before with 800 units of human chorionic gonadotropin (HCG, Lee BioSolutions). Collected eggs were fertilized in vitro using the sperm suspension described above and the developing embryos were cultured in 0.1x MMR and staged according to Nieuwkoop and Faber (8). For *in vitro* mRNA synthesis, plasmid DNA from pCS2-hTFG-3xFlag, pCS2-xTFG-3xFlag, pCS2-xDkk1, pCS2-nLacZ, pCS2-Lrp6-APEX2 and pCS2-xNotum were linearized with NotI and transcribed with SP6 RNA polymerase using an Ambion mMessage mMachine kit. The amount of mRNAs injected per embryo is indicated in the figure legends. For knock-down experiments, a TFG translation-blocking Morpholino (MO) antisense sequence recognizing both *Xenopus* homeolog alleles was designed and synthesized by Gene Tools: 5’ TTAAGTCCAGTTGTCCATTCATTGT 3’. 36 ng of TFG MO (1:1 dilution of the 1 mM stock solution) were injected per embryo into the 2 dorsal animal blastomeres at the 4 to 8 cell stage. Injection at the 8-cell stage resulted in a stronger phenotype. For Sox2 staining and the *pCSKA-xWnt8* DNA injection (43) experiment, embryos were injected 4 times (72 ng in total of TFG-MO) at the 2-cell (Sox2) or 4-cell stage (xWnt8 DNA, 32 pg total). A mixture of Bighead.L and Bighead.S MO (32 ng per embryo) was injected two times into the two dorsal blastomeres at the 4-cell stage, with or without TFG MO. Maternal knock-down of xLrp6 and host transfer of Lrp6-depleted oocytes was performed as previously described, using albino female frogs as hosts (9). LacZ lineage tracing and *in situ* hybridization using antisense probes for *Sox2, Engrailed-2, Krox20, TFG, Otx2, Rx2a* and *MyoD* were performed according to standard protocols (http://www.hhmi.ucla.edu/derobertis/). The *Xenopus* TFG *in situ* hybridization probe was generated by PCR using a reverse primer containing a T3 promoter (Table S1). The PCR product was then purified with a PureLink purification kit (Invitrogen^®^) and used as a template to synthesize the antisense probe using the T3 promoter, as previously described (10).

### Western Blots

Standard protocols were used to perform sodium dodecyl sulfate polyacrylamide gel electrophoresis (SDS-PAGE) and western Blot. A list of primary and secondary antibodies used in this study, including their dilutions, is provided in Table S2. Nitrocellulose membranes were blocked in TBST-milk 5%. Primary and secondary antibodies were diluted in TBST-milk 2.5%. For pLrp6 (Tp1479) western blot, nitrocellulose membranes were blocked with Li-COR blocking buffer (TBS-based) and Tp1479 antibody was diluted in the same blocking buffer supplemented with 0.2% Tween-20. Li-COR Odyssey 3.0 software was used to analyze the blotted membranes.

### Heatmaps

Heatmaps were generated in R-Studio (1.2.1335). For Fig. 2*C* and *SI Appendix*, Fig. S2*F*, normalized enrichment values of Wnt3a treated samples were used as inputs. Horizontal row z-scores were obtained from the values by calculating the mean, variance and standard deviation of each protein by using the gplots v3.0.1 package in R-Studio. The rows/proteins were left unclustered, as were the columns/treatments. The heatmap script used to generate our heatmaps was as follows:

##: Load gplots package

library(gplots)

##: Import Wnt dataset

Normalized.heatmap.of.Wnt.targets. <-read.csv(“∼/Desktop/APEX2 normalized data/Normalized heatmap of Wnt targets .csv”, row.names=1)

##: Convert data into a matrix frame

Normalized.heatmap.of.Wnt.targets._matrix <-data.matrix(normalized.heatmap.of.Wnt.targets.)

\## Generate heatmap via heatmap.2

heatmap.2(Normalized.heatmap.of.Wnt.targets._matrix, trace = “none”, scale = “row”, col = greenred(90), cexCol = 1, Colv = FALSE, Rowv = FALSE)

### Immunofluorescence

HeLa cells were grown on 12-well plates containing glass coverslips. For HEK293T cell immunostaining, coverslips were pre-coated with a solution containing 0.01% poly-L-Ornithine (Millipore) overnight at 37°C. Two days following DNA transfection, the cell culture medium was replaced with control or Wnt3a conditioned medium as required, and cultured as indicated in the figure legends at 37°C. Cells were washed twice with PBS (phosphate buffered saline, 137 mM NaCl, 2.7 mM KCl, 10 mM Na_2_HPO_4_, 1.8 mM KH_2_PO_4_), fixed in 4% (wt/vol) paraformaldehyde (Sigma, P6148) in PBS for 20 min, then permeabilized with 0.2% (vol/vol) Triton X-100 in PBS. Note that for cell-surface staining of Lrp6 in some experiments we used a lower concentration of Triton X-100 (0.05%), which markedly improved plasma membrane immunofluorescence (e.g., Fig. 1*F*). Coverslips were then washed with PBS, blocked for 1 hour in blocking buffer consisting of 3% (wt/vol) BSA in PBS at room temperature, and incubated with primary antibodies in blocking buffer overnight at 4°C. The next day cells were washed 3 times with PBS, incubated with secondary antibodies diluted in blocking buffer for 1 hour at room temperature and mounted onto glass slides with ProLong Gold antifade reagent with DAPI (Life Technologies) to stain cell nuclei. A list of primary and secondary antibodies used in this study, including their dilution, is provided in Table S3. Images were acquired using a Zeiss Imager Z.1 microscope with Apotome and 63X oil immersion objective. Image analysis was achieved with Zeiss (Zen) and ImageJ (NIH) imaging software.

### Image Quantification and Statistical Analyses

For quantitatively analyzing the Wnt3a-dependent increase in Hrs-containing vesicles, Hrs immunofluorescence was quantified in control and Wnt3a-treated cells using ImageJ software analyses (at least n=10 cells per condition). Briefly, this included normalizing fluorescence in images and measuring fluorescence in individual cells. Pearson’s correlation coefficients were calculated using ImageJ software to assess the degree of colocalization between puncta in two different channels. Two-tailed t tests were used for two-sided comparisons, and *P < 0.05, **P < 0.01, and ***P < 0.001 were considered to be statistically significant.

### Movies

Time-lapse movies were acquired with a Zeiss Observer.Z1 microscope equipped with Apotome.2. The microscope has a fully automated stage and a Temperature/CO_2_ Module S for cell culture. Images were collected using Colibri LED using green fluorescence filters with Apotome and a 63x LD (Long working Distance) Apochromat objective. Cells were grown on 4-well Nunc Lab-Tek II Chamber SlideTM treated with Fibronectin. Each chamber was incubated with 0.1 ml of 100 μg/ml of sterile Fibronectin (Sigma) for 30 min at room temperature, and washed 3 times with PBS before seeding cells in DMEM medium containing 10% Fetal Bovine Serum. Image acquisition was 40 frames in 20 min. Fluorescence filters were controlled by Axiovision 4.8 software and saved in this program. Movies were then processed using Adobe Premiere Pro CC 2019.

### Luciferase assays

To test Wnt3a conditioned medium (*SI Appendix*, Fig. S1*H* and Fig. S2*D*), a HEK293T cell line permanently transfected with β-Catenin-activated (BAR) reporter (11) Luciferase/Renilla system was used. Cells were plated in 12-well culture plates and incubated until the next day (at 60% of confluency) with either control or Wnt3a conditioned medium for 16 hours. After treatment, cells were lysed in 100 µl of Passive Lysis Buffer (Promega). Luciferase assays were performed with the Dual-Luciferase Reporter Assay System (Promega) using a Glomax Luminometer (Promega) according to manufacturer’s instructions. Renilla readings were used for normalization. For the siRNA knock-down experiment shown in Fig. 3*I*, the following procedure was adopted. On day one, 2 million HEK-293T cells were plated in 6-well culture dishes. On day two, cells were transfected with either siRNA targeting TFG (Dharmacon) or Scrambled sequences, using BioT transfection reagent, in triplicate. On day three, cells were re-plated onto 12-well plates. On day four, cells were transfected with BAR-Luc Wnt reporter DNA, pCS2+ Renilla and pCS2+ carrier DNA, with each well receiving a total of 2 μg of DNA. The following DNA amounts were used: 1.2 μg BAR-Luc reporter, 0.4 μg pCS2+ Renilla, and 0.4 μg of pCS2+ empty vector. 24 hours following DNA transfection, cells were incubated with either Wnt3a or control medium for 16 hours, and Wnt activation analyzed by Luciferase assays as described above. For the luciferase assays shown in Figure 4, WT or TFG KO HEK293T BR (BAR/Renilla) reporter cells were treated with control or Wnt3a conditioned medium, or CHIR99021 5 µM final concentration, for 16 hours at 37 °C. Cells were then lysed and analyzed by luciferase assay as described above.

### RT-PCR on *Xenopus* cDNA

*Xenopus* embryos were collected at the desired developmental stages, and total RNA was extracted from groups of 3 embryos each with the RNeasy Mini Kit (QIAGEN) following manufacturer’s instructions. 350 µl of lysis buffer were used per group of embryos, and on-column DNAse treatment was performed to remove genomic DNA following the Qiagen protocol. 2 µg of total RNA was used for cDNA synthesis with random hexamer primers, using the AffinityScript Multi-Temp Reverse Transcriptase (Agilent). cDNA was then used for semiquantitative RT-PCR, and the PCR products were resolved on 0.8% Agarose gel. Primer sequences are listed in Table 1.

